# Inhibitor binding influences the protonation states of histidines in SARS-CoV-2 main protease

**DOI:** 10.1101/2020.09.07.286344

**Authors:** Anna Pavlova, Diane L. Lynch, Isabella Daidone, Laura Zanetti-Polzi, Micholas Dean Smith, Chris Chipot, Daniel W. Kneller, Andrey Kovalevsky, Leighton Coates, Andrei A. Golosov, Callum J. Dickson, Camilo Velez-Vega, José S. Duca, Josh V. Vermaas, Yui Tik Pang, Atanu Acharya, Jerry M. Parks, Jeremy C. Smith, James C. Gumbart

## Abstract

The main protease (M^pro^) of severe acute respiratory syndrome coronavirus 2 (SARS-CoV-2) is an attractive target for antiviral therapeutics. Recently, many high-resolution apo and inhibitor-bound structures of M^pro^, a cysteine protease, have been determined, facilitating structure-based drug design. M^pro^ plays a central role in the viral life cycle by catalyzing the cleavage of SARS-CoV-2 polyproteins. In addition to the catalytic dyad His41-Cys145, M^pro^ contains multiple histidines including His163, His164, and His172. The protonation states of these histidines and the catalytic nucleophile Cys145 have been debated in previous studies of SARS-CoV M^pro^, but have yet to be investigated for SARS-CoV-2. In this work we have used molecular dynamics simulations to determine the structural stability of SARS-CoV-2 M^pro^ as a function of the protonation assignments for these residues. We simulated both the apo and inhibitor-bound enzyme and found that the conformational stability of the binding site, bound inhibitors, and the hydrogen bond networks of M^pro^ are highly sensitive to these assignments. Additionally, the two inhibitors studied, the peptidomimetic N3 and an *α*-ketoamide, display distinct His41/His164 protonation-state-dependent stabilities. While the apo and the N3-bound systems favored N_*δ*_ (HD) and N_*ϵ*_ (HE) protonation of His41 and His164, respectively, the *α*-ketoamide was not stably bound in this state. Our results illustrate the importance of using appropriate histidine protonation states to accurately model the structure and dynamics of SARS-CoV-2 M^pro^ in both the apo and inhibitor-bound states, a necessary prerequisite for drug-design efforts.

## Introduction

A new coronavirus, named severe acute respiratory syndrome coronavirus 2 (SARS-CoV-2), has caused a worldwide outbreak of viral pneumonia starting in December 2019, known as coronavirus disease 2019 (COVID-19), and, currently, treatment options are very limited. Two potentially druggable targets in SARS-CoV-2 and other betacoronaviruses are the chymotrypsin-like protease (3CL M^pro^) and the papain-like protease (PL^pro^), which are responsible for cleaving the large polyproteins translated from the viral RNA. M^pro^ has been shown to cleave at least 11 sites of the polyprotein 1ab, recognizing the sequence Leu-Gln↓Ser/Ala, typically denoted P2-P1↓P1’, with the cleavage occurring between the P1 and P1’ residues, as denoted by the arrow.^1–3^ Because this sequence is not recognized by any known human protease, M^pro^ is an attractive target for SARS-CoV-2-specific antivirals, and previous attempts at developing them for other coronaviruses focused on it.^1,4–9^

Crystal structures of M^pro^ from SARS-CoV, SARS-CoV-2, and other coronaviruses reveal that this enzyme is a homodimer and is structurally conserved among coronaviruses.^7,8,10,11^ M^pro^ is a cysteine protease that employs a non-canonical Cys-His catalytic dyad with the overall structure of the M^pro^ dimer illustrated in Figure 1. The individual monomer consists of three domains: domain I (residues 8-101), domain II (residues 102-184), and domain III (residues 201-303); domains I and II interact to create the catalytic site, while the *α*-helical domain III is responsible for dimerization. ^7,8^ Given that earlier work has shown that the M^pro^ monomer is catalytically inactive,^12–14^ it appears that dimerization is essential for M^pro^ catalytic activity. The first residue, Ser1, of the N-finger (residues 1-7) interacts with Glu166 of the other monomer, shaping and stabilizing part of the substrate binding site.^12^ The dimerization is abolished by removal of the N-finger and mutation of Tyr161, while the dimerization binding constant is reduced for E166A mutants.^12,14,15^

**Figure 1:**
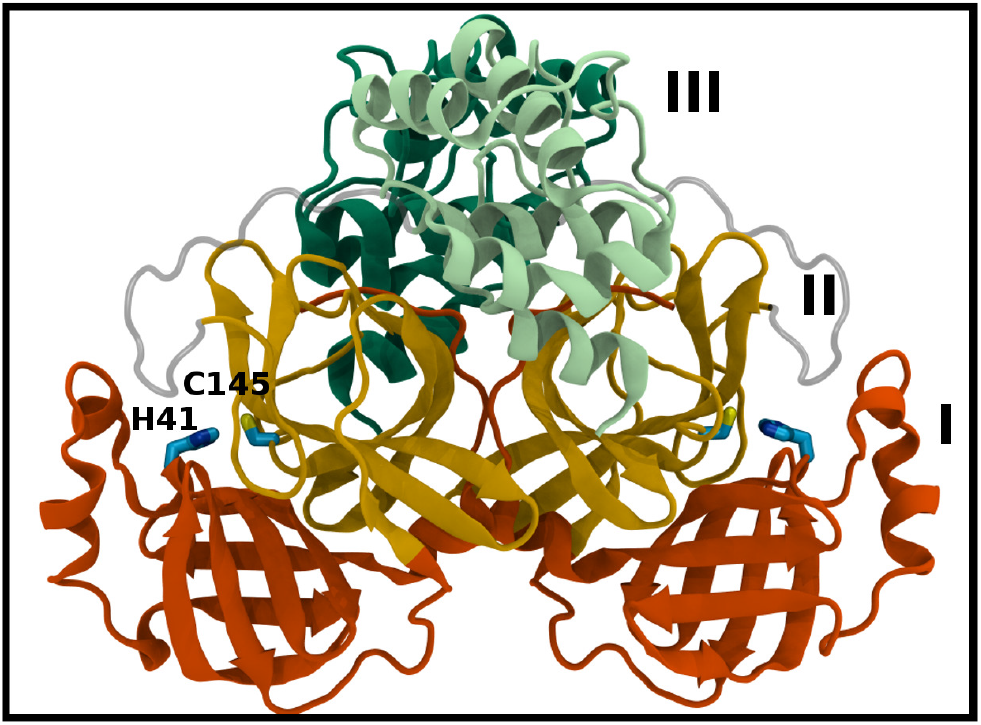
M^pro^ dimer structure and binding site interactions (PDB entry 6WQF). M^pro^ homodimer with the three domains illustrated and color coded as follows: Domain I (dark orange), domain II (gold), and domain III (light green/dark green monomer A/B) with the catalytic dyad residues, His41 and Cys145 (rendered in licorice).

Given the urgent need for effective treatment options for COVID-19, the search for M^pro^ inhibitors has been intense. For example, several inhibitors have been identified with crystal structures obtained and released, including the peptidomimetic N3 and *α*-ketoamides (Figure S1). ^7,8^ These inhibitor-bound structures provide excellent starting points for further drug optimization strategies. In addition to the crystal structures of M^pro^, both in its apo as well as ligand-bound states,^7,8,11^ recent crystallographic screening^16^ has revealed the binding of a multitude of molecular fragments both in the active site as well as at the dimer interface where oligomerization can be disrupted. Such structural studies, complemented by computational tools for drug design, offer tremendous promise for the rapid generation of potent lead compounds for the production of effective antivirals.

M^pro^ of coronaviruses employs a His-Cys catalytic dyad for protein cleavage. Mutation of the cysteine to serine reduces catalytic efficiency tenfold, and the enzyme is most active at pH 7.4. ^14,17,18^ As observed in neutron crystallographic structures of other proteins, histidines can form hydrogen bonds with main-chain amide nitrogens, resulting in low pKas. ^19,20^ Such histidines will remain in a neutral deprotonated state; for example, in M^pro^ these include His172 and His246. Several other histidines are positioned near the binding site: His41 (catalytic), His64, His80, His163, His164 and His172, and it has been suggested that protonation and deprotonation of some of these histidines are responsible for changes in enzymatic activity in SARS-CoV.^17,21^ Specifically, crystal structures obtained at pH 6.0 suggest that His163 becomes protonated, causing rearrangement of interactions with the N-finger and partial collapse of the binding site.^17^ These observations were supported by molecular dynamics (MD) simulations.^21^ It was also suggested that in SARS-CoV M^pro^, His172 is protonated because both nitrogens of its side chain appear to donate hydrogen bonds (PDB entries 1UK3, 2DUC).^17^ For the same protein, MD simulations showed that protonation of His172 increased the flexibility of Glu166 and in turn increased the distance between Glu166 and the N-finger.^21^ However, it is not clear how Glu166 flexibility affects enzyme catalysis. Furthermore, released structures of SARS-CoV-2 M^pro^ show greater agreement between the two monomers than those of SARS-CoV M^pro^ and none of them show His172 potentially donating two hydrogen bonds.^7,8^ Therefore, to determine the role of the protonation state of His172 in SARS-CoV-2 M^pro^, further investigation is needed.

Since the release of SARS-CoV-2 M^pro^ structures in apo and inhibitor-bound states, multiple MD simulation and docking studies of this enzyme have already been published.^22–28^ Although a precise determination of the specific protonation states of Cys145 and several histidines in M^pro^ in this pH-sensitive enzyme will be critical to effective and robust computational drug design efforts, these protonation states have not been unequivocally determined. Despite the fact that X-ray crystallography can identify and precisely position non-hydrogen atoms, the low electron density of hydrogen normally precludes the direct X-ray determination of protonation states. Neutron crystallography, on the other hand, can often identify the location of hydrogen atoms even at modest resolution.^29–31^ However, protonation states in proteins can deviate considerably from intrinsic (i.e., aqueous solution) values due to preferential stabilization of protonated or deprotonated states by interactions with ligands and the protein interior. Furthermore, altering protonation states of titratable groups can modulate protein dynamics and stability via dynamic hydrogen-bonding networks, as has recently been illustrated using a novel graph-based analysis approach for the SARS-CoV-2 S protein. ^32,33^

The protonation states of the catalytic residues, His41 and Cys145, and His164 in SARS-CoV M^pro^ have also been investigated previously.^18,34^ Mechanistic studies concluded that the catalytic residues are most likely neutral in the active state,^18^ while MD simulations showed increased distances between the catalytic dyad atoms NE of His41 and S of Cys145 in the charged state in comparison to the neutral state. ^34^ However, combined protonation of His164 and deprotonation of Cys145 could not be excluded.^34^ In addition, it was shown that a water molecule actively participates in the cleavage reaction.^34–36^

Therefore, given similar ambiguity in the protonation states of SARS-CoV-2 M^pro^, we have investigated the stability of 12 possible protonation states of this protein. MD simulations of apo and inhibitor-bound M^pro^ dimers were performed for each state and the resulting structural and dynamical properties were analyzed. Two inhibitors were studied: N3 and one of the most potent *α*-ketoamides to date (Figure S1), which we refer to simply as ketoamide hereafter. Overall, we conclude that the structural stability of M^pro^ is indeed affected by the assignment of His protonation states in simulation studies, and the most structurally stable protonation states vary in a ligand-dependent manner.

## Methods

### Simulations

All simulations used the CHARMM36m force field for proteins and the CHARMM-modified TIP3P water model.^37,38^ Initial CGenFF parameters for the inhibitors N3 and ketoamide were taken from the CGenFF program. ^39–41^ Ketoamide charges and bonded parameters that had poor analogies according to the CGenFF scoring system were reoptimized using the force field toolkit (ffTK);^42^ see the supplementary information (SI) for details. Gaussian16 was used for QM calculations required for parameter optimization. ^43^ N3 parameters were not further optimized because they had lower penalties from CGenFF than ketoamide, and because those penalties mainly involved the methylisoxazole ring, which does not interact with M^pro^ in the crystal structure.

NAMD 2.13 was used for a set of 20-ns simulations of each potential protonation-state combination. ^44,45^ All covalent bonds with hydrogens were kept rigid, allowing integration of the equations of motion with a 2-fs time step. A 12 Å cutoff was used for van der Waals interactions and a smoothing function was applied from 10-12 ^Å^, ensuring a smooth decay to zero. The particle-mesh Ewald method was used to account for long-range electrostatic interactions. ^46^ Pressure and temperature were kept constant at biologically relevant values of 1 bar and 310 K using a Langevin thermostat and barostat, respectively. ^47^

The shorter simulations exploring alternative protonation states were run for 20 ns × 3 runs for all 12 states for two apo structures, one collected at 100 K (PDB entry 6YB7)^48^ and one collected at room temperature (PDB entry 6WQF),^11^ as well as for the ketoamide and N3-bound structures (6Y2G^7^ and 7BQY,^8^ respectively), giving a cumulative total of 2.88 *μ*s simulation time. Both 6Y2G and 7BQY are missing the positions of C-terminal amino acids 302-306. Hence, the coordinates for these residues were obtained from two other M^pro^ structures with intact C-termini. We used the older N3-bound structure (PDB entry 6LU7) for the C-terminus of 7BQY and the apo structure 6YB7 for the C-terminus of 6Y2G. Although the inhibitors are covalently bound in both crystal structures, we chose to simulate them in their pre-covalent states, prior to bond formation. Previous QM/MM studies have investigated the pre-covalent states using the covalently-bound structures in order to study the cleavage mechanism. ^34,36^ Here we used pre-covalent states because we are predominantly interested in the effects of the protonation states on non-covalent binding, which may be masked by covalent attachment. All structures simulated with NAMD were solvated and ionized with 0.15 M NaCl in VMD. ^49^ After minimization, water and ions were equilibrated for 1 ns while the protein and inhibitor, if present, were restrained with a force constant of 2 kcal/mol/Å2. In a second equilibration step only the protein backbone was restrained with the same force constant for 4 ns. For analysis, the first 5 ns of the subsequent unrestrained 20-ns simulation runs was discarded.

#### Extended simulations

The extended simulations focusing on a more limited set of protonation states, in which His41 from 6WQF model was assigned as either HD41 or HE41 paired with HE164, were simulated using Gromacs 2020.1,^50^ with five sets of 250-ns simulations run to permit analysis over longer time scales. The molecular systems were built directly from the initial crystal structures, with additional solvent and neutralizing ions added with GROMACS tools. For consistency with the CHARMM36m force field used throughout this study, ^38^ the same 12 Å cutoff was applied, with particle-mesh Ewald^46^ used for long-range electrostatics. To maintain the temperature at 310 K, a velocity rescaling thermostat was used, with an isotropic Parrinello-Rahman barostat maintaining the pressure at 1 bar. ^51^ LINCS was used to restrain the lengths of bonds to hydrogen atoms, enabling 2-fs time steps. ^52^

#### Free-energy perturbation

The relative protein-ligand binding free energies associated to the transformations of residue HD41 into HE41, on the one hand, and residue HE163 into HP163, on the other hand, with either the ketoamide or N3 bound to the enzyme, were determined using free-energy perturbation (FEP).^53,54^ Towards this end, the protonation state changes were carried out in the apo (unbound) state of the enzyme and in the respective inhibitor-bound state, concomitantly in both monomers of the homodimeric protein. The following protonation states were used for the residues that were not altered in the FEP transformation: HD41, HE164, HE163, HE172 and neutral C145. For both His41 and His172, the transformations were first initiated from the HE state. The change in free energy ΔG° was obtained from the difference in alchemical free energies between the bound and unbound states, i.e., 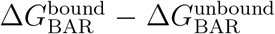. Considering the nature of the alchemical transformations, the reaction path was stratified into 100 stages of equal widths. Replacement of HD41 and HE163 was performed over 160 ns, both in the bound and unbound states (see Table S4 for individual runs). The dual-topology paradigm was utilized,^55^ whereby the initial and the final states of the alchemical transformation coexist, albeit not interacting. To improve sampling efficiency, geometric restraints were introduced to ensure that the imidazole rings of the alternate topologies remain superimposed in the course of the amino-acid replacement. At each stage of the stratified reaction path, data collection was prefaced by suitable thermalization in the amount of one-fourth of the total sampling. To augment the accuracy of the free-energy calculation, each alchemical transformation was carried out bidirectionally,^56^ and the associated relative binding free energy was determined using the Bennett acceptance ratio (BAR) method.^57^ The error bars associated to the relative free energies were computed from the hysteresis between the forward and backward alchemical transformations of the bidirectional FEP calculations, and assume independence of the transformations in the bound and unbound states. All free-energy calculations were performed with the recent single-node path implementation of FEP^58^ available in NAMD 3.0.^45^

### Analysis

Root-mean-square deviation (RMSD) of the proteins was measured by comparing the C_*α*_ positions to those of the starting structures. This property was calculated for each monomer separately after alignment and aimed at detecting differences in the overall protein structure. The RMSD of the active site was calculated using all non-hydrogen atoms for residues 25 to 28, 38-50, 139-145, 160-176, and 185-195 of the selected monomer and residues 1 and 2 of the other monomer (Figure S3A). We aimed to investigate the changes in the shape of the inhibitor binding site by analyzing this property. Therefore, the RMSD of the site was calculated after alignment of the above-defined active site residues with their positions in the starting structures. The RMSD of the inhibitor was measured by comparing its position over time to its crystallographic position after aligning to the protein. Specifically, this RMSD was calculated after aligning to C_*α*_ positions of the dimer complex. All properties were calculated separately for each of the two monomers.

All distribution plots were generated by applying Gaussian kernel density estimation on normalized histograms using the seaborn python package. This estimation was used to generate smoother curves, which are easier to compare and interpret.

Hydrogen bonding analysis was performed with the hydrogen bonding plugin of VMD 1.9.4 with a 3.5-Å and 35-degree heavy-atom distance and angle cutoffs, respectively. Hydrophobic contacts were straightforwardly determined by counting the number of times each ligand carbon atom comes within 4.5 A of a protein carbon atom. Pocket volume was computed using the Epock VMD plugin with the standard program settings. ^59^ See the SI for definitions and visualization of the binding sites and volumes (Figure S4).

## Results and Discussion

The structure of the substrate binding site for the main protease of SARS-CoV-2 includes the catalytic dyad residues (Cys145, His41), as well as recognition pockets for specific substrate residues. These include the S1 pocket, providing His163 for interaction with the highly conserved substrate residue glutamine, and the S2 pocket, which accommodates the hydrophobic P2 leucine. These subsites are often the target for antiviral drug design, and, depending on the length of the inhibitor, additional subsites may be occupied. For example, the long peptidomimetic inhibitor N3 occupies the S1, S2, S4, and S5 sites and extends into the S1’ site.^8^

We investigated the effects on the structural properties of the apo and ligand-bound systems by altering the protonation state of the catalytic dyad, Cys145 and His41, as well as those of three histidines near the substrate binding site, His163, His164, and His172. Cys145, His41, and His163 form direct contacts with substrates and inhibitors. His163 HD also forms a hydrogen bond with Tyr161, while His41 is also involved in a network of interactions that includes a water molecule, His164, and Asp187, which in turn is stabilized by a salt bridge with Arg40. In addition, NE of His164 forms a hydrogen bond with the side chain of Thr175. In the structure 6YB7, a hydrogen bond between the His41 side chain and the Gly174 backbone is observed, which is absent in the other three structures studied here. The Arg40-Asp187 interaction bridges domain I with the interface of domains II and III. It has been suggested that the water molecule acts as a third partner in the enzymatic activity of M^pro^.^11,34^ Many successful inhibitors^7,8^ of M^pro^ include a lactam moiety, mimicking the glutamine of the substrate, which binds in the S1 substrate subsite via a hydrogen bond with His163. Maintenance of the S1 pocket^7^ and dimerization state^14,15,60^ have been shown to affect the catalytic efficiency of M^pro^. In addition to His163, residues implicated in maintaining either the shape of the S1 pocket or the dimerization state of the protein include Glu166, His172, Phe140 and the N-terminal serine, Ser1’, from the other monomer (Figure 2A). For both N3 and ketoamide, a hydrophobic moiety occupies the S2 subsite, which is a relatively deep pocket formed by Met49, Tyr54, Met165, as well as the hydrocarbon portion of the Asp187 side chain.^7,8^ These regions are illustrated in Figure 2B and 2C for the N3 and ketoamide-bound structures, respectively.

**Figure 2:**
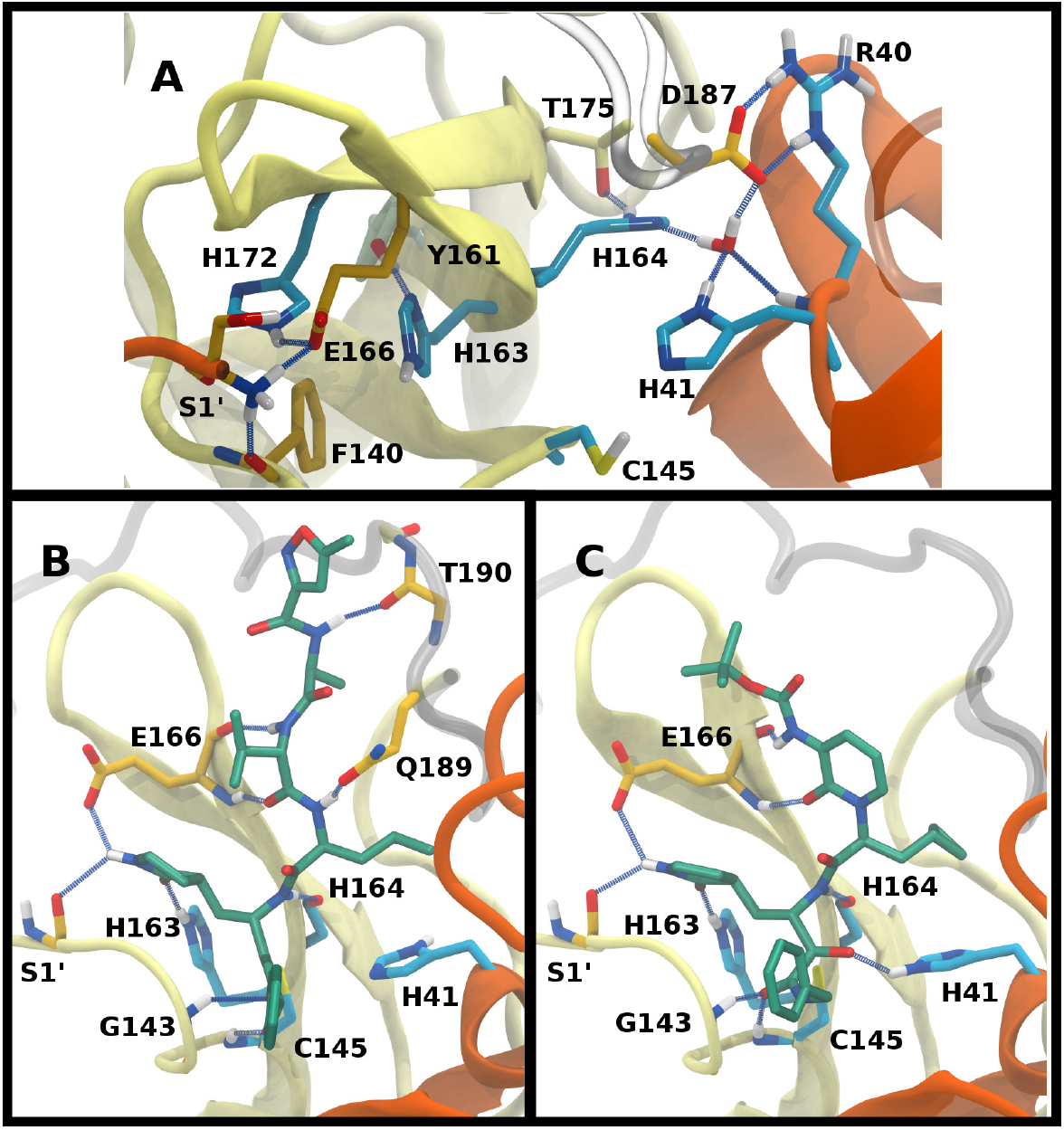
Hydrogen bonding interactions in the catalytic site for the apo, N3-bound, and ketoamide-bound structures. A) Illustrated for the HD41-HE164 apo state taken from the simulation are the i) catalytic dyad, ii) the crystallographic water bridging His41, His164, and Asp187 as well as His164 with Thr175, and iii) S1 pocket interactions. The N-terminal serine residue is labeled with a prime to indicate that it is from the alternate monomer. B) Binding pocket of N3-bound M^pro^ with hydrogen bonds displayed taken from the HD41-HE164 state. C) Binding pocket of ketoamide-bound M^pro^ with hydrogen bonds displayed, taken from the HE41-HD164 state. The ligands in both B) and C) are rendered in licorice with carbon atoms in green. Hydrogen bonds are illustrated with blue lines.

### Setup of the studied systems

Various methods allow for computation for pKas for specific residues. Most straightforward are webservers, e.g., the Poisson-Boltzmann solver H++^61^ and the empirical predictor propKa. ^62^ Importantly, however, while the former predicts HD and HE states of histidine, the latter only distinguishes between protonated and neutral states. Nonetheless, the assignment of histidine protonation states for subsequent detailed studies, e.g., virtual screening, remains challenging. ^63^ A more accurate, albeit computationally costly approach, is constant pH molecular dynamics (CpHMD), which was recently applied to PL^pro^ of SARS-CoV-2.^64,65^ Although CpHMD can reveal likely combinations of protonation states, we also wanted to consider the structural consequences of unexpected ones. Therefore, shorter, repeated standard MD simulations with fixed protonation states were used.

The following naming scheme was used for different protonation states: C/CD for neutral/deprotonated Cys145 and HE/HD/HP for histidine protonated on NE, ND, or both nitrogens, respectively. The protonation states of His64, His80 and His246 were not changed from their assigned HE, HD and HE states, respectively, because alternative protonation states were either unlikely or unimportant. Specifically, His64 is solvent exposed, His80 ND forms a hydrogen bond with side-chain oxygen of Asn63, and His246 ND forms a hydrogen bond with backbone NH of Thr243. In addition, the HD state of His163 was not considered because it would prevent hydrogen bonding with the substrates and several known inhibitors.^7,8^ Because simultaneous protonation of ND nitrogens in His41 and His164 was expected to cause sterical repulsion based on available crystal structures we only included one state with simultaneous ND protonation of both residues.^7,8,11^ Table 1 shows the 12 protonation states that were investigated. The protonation states of Cys145, His41 and His164 were considered to be interdependent because of their proximity, while the protonation states of His163 and His172 were assumed to be independent from the other residues. The simulations of these states were performed for two apo crystal structures and two inhibitor bound structures, one with N3 and one with a ketoamide (see Methods for details).

**Table 1:** Histidine protonation state combinations considered in this work. Altered protonation states are shown in bold. The corresponding groups and names of the states are also shown.

| Cys145 | His41 | His163 | His164 | His172 | Name | Group |
| --- | --- | --- | --- | --- | --- | --- |
| C | **HE** | HE | **HD** | HE | HE41-HD164 | 1,2,3 |
| **CD** | **HP** | HE | **HD** | HE | CD145-HP41-HD164 | 1 |
| **CD** | **HE** | HE | **HP** | HE | CD145-HE41-HP164 | 1 |
| C | **HP** | HE | **HE** | HE | HP41-HE164 | 2 |
| C | **HE** | HE | **HE** | HE | HE41-HE164 | 2 |
| C | **HE** | HE | **HP** | HE | HE41-HP164 | 2 |
| C | **HD** | HE | **HD** | HE | HD41-HD164 | 2 |
| C | **HD** | HE | **HE** | HE | HD41-HE164 | 2 |
| C | **HE** | **HP** | **HD** | HE | HE41-HP163-HD164 | 3 |
| C | **HE** | HE | **HD** | **HD** | HE41-HD164-HD172 | 3 |
| C | **HE** | HE | **HD** | **HP** | HE41-HD164-HP172 | 3 |
| C | **HD** | **HP** | **HE** | HE | HD41-HP163-HE164 | 3 |

To facilitate the analysis, the protonation states were divided into three groups, labeled Group 1, 2 and 3, with the goal of determining the most favorable protonation state of Cys145, His41-His164, and His163-His172, respectively. To compare the stability of systems with different protonation states, we computed several properties, namely root-mean-square deviation (RMSD); root-mean-square fluctuations (RMSF); hydrogen bonding; hydrophobic contacts; water occupancy near His41, His164, and Asp187; and volume of the binding pocket. The RMSD was calculated for the C_*α*_ atoms of the full protein, for the binding site alone, and for the inhibitor relative to the protein. Hydrogen bonding was measured for relevant residues of the protein and between the bound inhibitors and the protein. Residue pairs monitored include the catalytic dyad (Cys145 and His41), those implicated in maintaining the shape of the S1 specificity pocket (His163-Tyr161, His172-Glu166, and the intermonomer interactions Ser1’-Glu166 and Ser1’-Phe140), and those in regions surrounding the putative catalytic water (Arg40-Asp187). Finally, given the hydrophobic moieties in the N3- and ketoamide-bound M^pro^ structures, particularly the leucine/cyclopropyl group that occupies the S2 subsite, hydrophobic contacts were also measured between the protein and inhibitor. Based on the analysis of these properties, the most structurally stable states were determined.

### Apo enzyme

Simulations of the two crystal structures of the apo enzyme state (PDB entries 6WQF and 6YB7, respectively^11,48^) were performed. Both structural models include full-length M^pro^, i.e., residues 1-306. The RMSDs, RMSFs, His41(NE) - Cys145(S) distances, and hydrogen bonding patterns are reported in Figures 3 and S5-S7 for the 6WQF simulations and in Figures S8-S11 for 6YB7.

**Figure 3:**
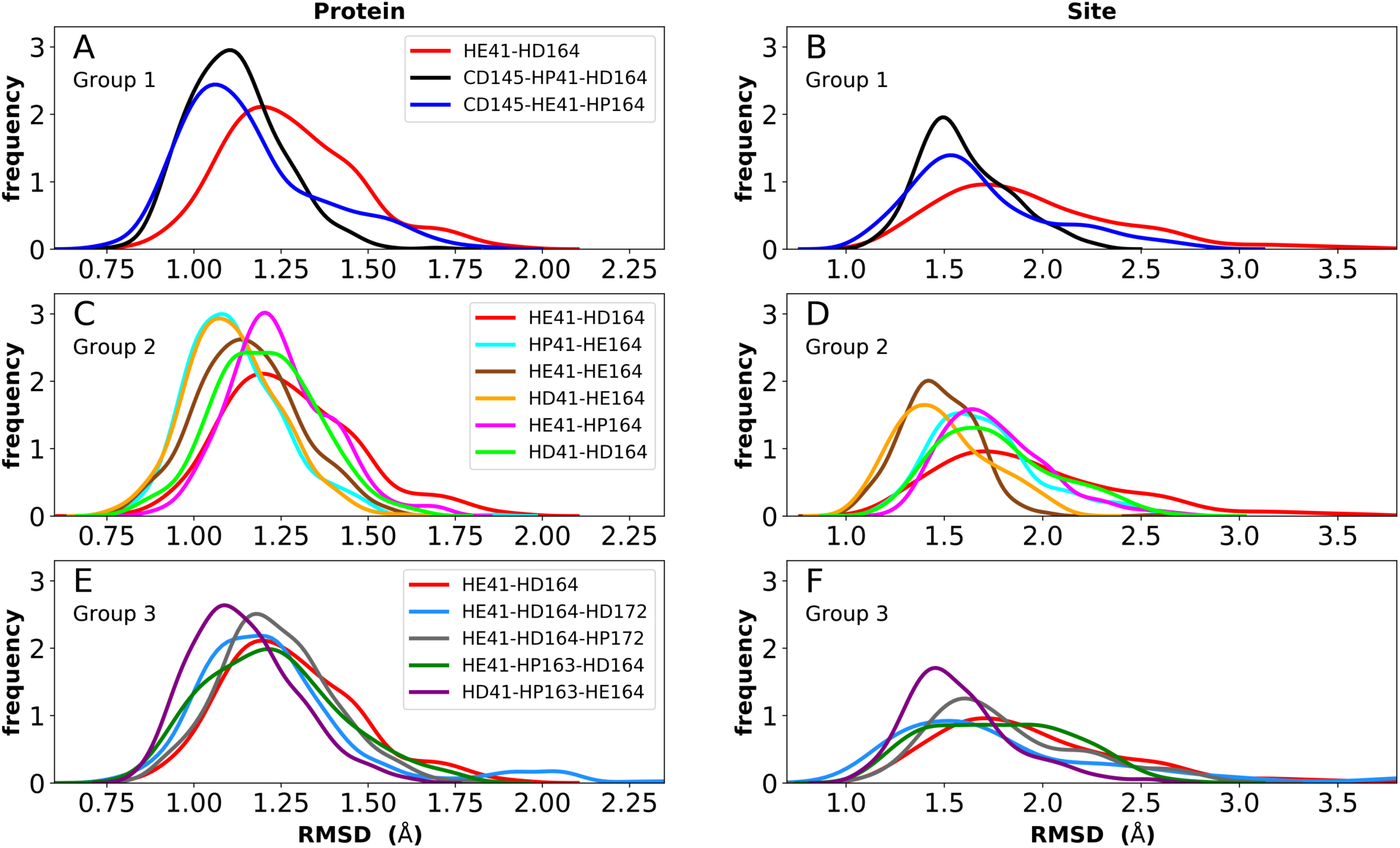
RMSD distributions for the three protonation-state groups from simulations of the apo structure (PDB entry 6WQF^11^). A-B) RMSD of (A) protein and (B) active site for Group 1. C-D) RMSD of (C) protein and (D) active site for Group 2. E-F) RMSD of (E) protein and (F) active site for Group 3.

For the 6WQF model, the following states had the lowest RMSDs for both the protein overall and the active site (Figure 3): CD145-HP41-HD164 (Group 1), HD41-HE164 (Group 2), HE41-HE164 (Group 2), and HD41-HP163-HE164 (Group 3). Notably, HE41-HD164-HP172 resulted in increased fluctuations of Ser1, suggesting instability of the N-finger in this state (Figure S5E) and decreased hydrogen bonding capacity (Figure S7C,E). The RMSF profiles (Figure S5) are similar for the different protonation states, and all states showed increased flexibility around residues 49-51 and 192-194, in agreement with previous simulations as well as B-factors for structure 6WQF. ^11^

To better differentiate between low-RMSD states, we evaluated the ability of previously identified residue pairs (Figure 2A) to form hydrogen bonds in each of the 12 protonation states starting from the apo structure (PDB entry 6WQF). Their occupancies are reported in Figure S7. Illustrated in Figures 4 and 5 are persistent hydrogen bonds as well as cases where perturbations of the S1 pocket interactions or hydrogen bonding near the crystallographic water occur. Although many of these interactions have similar propensities, particularly when considering the standard deviations, we used hydrogen bonding involving several specific interactions in the S1 pocket and the catalytic dyad (Figure S7) to assist in eliminating protonation states.

**Figure 4:**
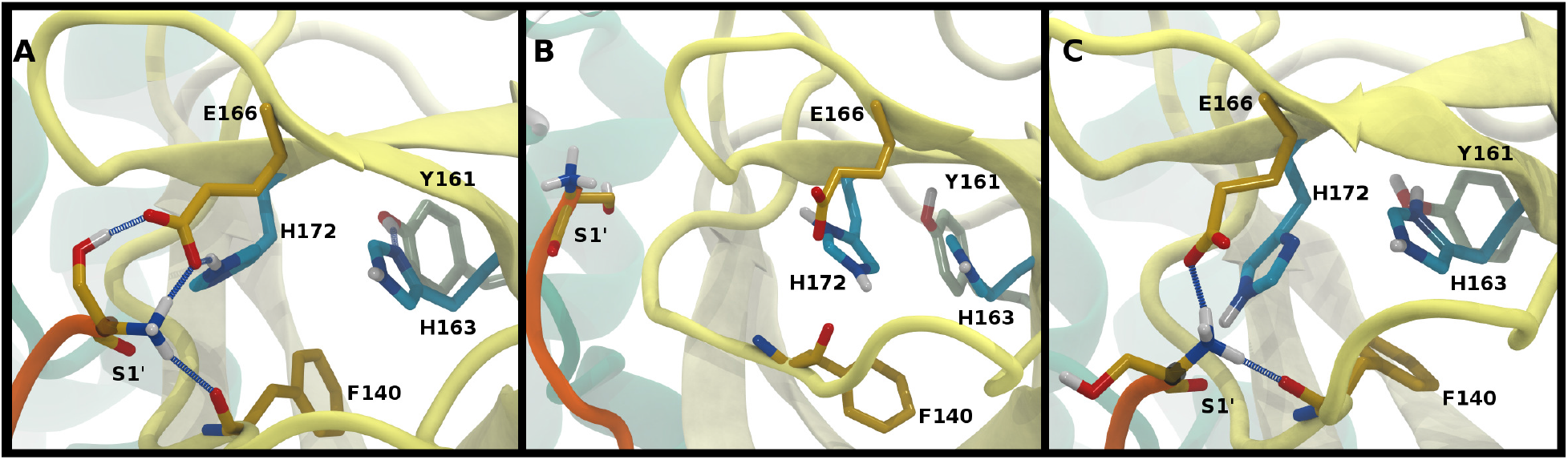
Hydrogen bonding in the S1 pocket. Example configurations of A) HD41-HE164 characteristic of robust S1 pocket interactions, B) HE41-HD164-HP172 illustrating the rupture of the S1’-Glu166 interaction and loss of the His163-Tyr161 hydrogen bond, and C) HE41-HP163-HD164 depicting the loss of the Tyr161 hydrogen bond donation and the His172-Glu166 interaction.

**Figure 5:**
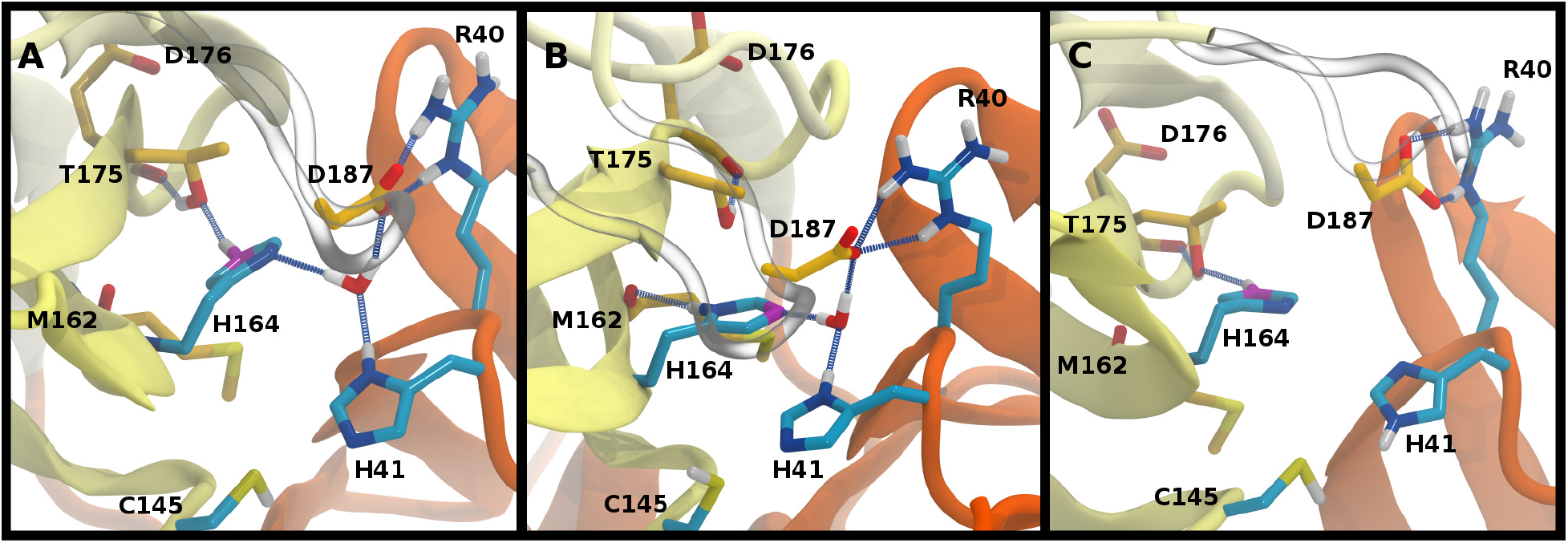
Hydrogen bonding involving the crystallographic water and His164 local environment. In each image, the NE nitrogen in the His164 side chain is colored magenta. A) HD41-HE164 illustrating hydrogen bond donation from His164 to Thr175. B) HD41-HD164 illustrating the His164 side chain rotation such that a hydrogen bond is formed with the backbone carbonyl of Met162. C) HE41-HE164 illustrating the disruption of the hydrogen bonding interactions and loss of the crystallographic water.

#### S1 specificity pocket interactions

For states with a neutral His163, the His163-Tyr161 side chains are engaged in a hydrogen bond with the histidine/tyrosine acting as the acceptor/donor, respectively (Figure 2A). This result is in agreement with known hydrogen bonding propensities for tyrosine residues. ^66^ The most dramatic differences occur upon protonation of His163 or His172 (Figures 4 and S7). For the single HP172 state considered, there is a significant reduction in the hydrogen-bonding occupancy for the His163-Tyr161 pair (Figures 4B and S7A). Moreover significant separation of the Ser1’ and Glu166 residues is observed, resulting in a disruption of this S1 pocket interaction (Figures 4B and S7C). This is consistent with the structural instability of the N-finger region in the HE41-HD164-HP172 state discussed above. When His163 is protonated (the HP163 states), the S1 pocket His163-Tyr161 interaction is ruptured due to loss of the hydrogen bonding acceptor moiety in the charged histidine side chain (Figures 4C and S7A). Only in the HD41-HP163-HE164 system does the tyrosine(acceptor)/histidine(donor) pair occur; however, the occupancy for this interaction (<20%) is quite low. Furthermore, significant perturbation of the hydrogen bonding between His172 and Glu166 is also observed in the charged HP163 states (Figures 4C and S7B). This finding is in accord with the suggestion by Tan et al.^21^ that protonation of His163 will contribute to altering the properties of the S1 specificity pocket. Given the disruption of the hydrogen bonding just discussed for HP163, the low-RMSD state from Group 3, HD41-HP163-HE164 may be discarded. Also of note is the observation that the S1 pocket interactions are significantly disturbed in the HD172 state (HE41-HD164-HD172; Figure S7) reinforcing the removal of this set of protonation states.

#### Catalytic Dyad (Cys145, His41)

The NE-S distance between the two catalytic residues is centered around 4 A for most states with neutral Cys145 and is slightly reduced for the states with deprotonated Cys145 (CD145) and/or charged HP41 (Figure S6B,D,F). Moreover, direct hydrogen bonding between the catalytic dyad residues Cys145 and His41 is rare, with non-zero occupancies only for the protonated systems, CD145-HE41-HD164 and CD145-HP41-HD164, at 22% and 26%, respectively (Figure S7F). These results are in general accord with the observed Cys145-His41 distance distributions discussed above and the elongated/weaker hydrogen bonding propensities of sulfur atoms. ^67^ Given the longer distances seen in the crystal structures and the shifted NE-S distances in the CD145/HP41 simulations we have removed the zwitterionic CD145-HP41-HD164 (Group 1) state from further consideration.

#### Crystallographic water

Interactions with water are particularly relevant for His41, which interacts with both His164 and Asp187 via a bridging crystallographic water (Figure 2A). This water molecule has been suggested to play a role in the catalytic reaction by stabilizing the positive charge accumulated on His41 and by assisting in proton transfer from Cys145 to His41.^11,34^ Given the possibility of water exchange, we have computed the occupancy of the putative catalytic water site by obtaining the fraction of frames in which a water molecule is simultaneously within 3.5 ^Å^A of His41(ND1), His164(ND1), and Asp187(OD1/OD2). The average occupancy is substantially above 80% for only the HD41-HE164 and HD41-HD164 states (Table S3), suggesting a particularly stable structural arrangement of His41, His164, Asp187, and water. It is noteworthy that in the HD41-HD164 state, the His164 changes conformation; the resulting geometry is analogous to that observed for the HD41-HE164 state. This similarity is clearly illustrated in Figures 5A and B, where the NE nitrogen on His164 has been colored magenta to highlight this rotation. This alteration of conformations produces the increased water occupancy observed in the HD41-HD164 state (Table S3) and is consistent with the increased site RMSD distribution observed for this system relative to HD41-HE164 (Figure 3D). Lastly, illustrated in Figure 5C is an example from the HE41-HE164 state, in which there is a local rearrangement of interactions with loss of the crystallographic water. In two out of the three 20-ns HE41-HE164 trajectories, this water is released into bulk.

Rotation of the side chain of His164 is characteristic of the HD states and generates additional state dependent hydrogen bonding propensities for this residue. We observe that in the HE and HP protonation states of His164 (Figure 5A), a hydrogen bond is present between the side chains of His164 (NE protonated and donating) and Thr175 (accepting). In contrast, for the His164 HD states the proton on the ND nitrogen is involved in a hydrogen bond with the backbone carbonyl of Met162 as illustrated in (Figure 5B). This occurs as a result of a rotation of the His164 side chain. Finally for 11 of the 12 states studied, the side chain of Thr175 donates a hydrogen bond to the main chain carbonyl of Asp176 with occupancy greater than 75% (Figure 5). The only exception is the HE41-HP163-HD164 state where the Thr175 side chain donates a hydrogen bond to HD164 (~50%).

#### Pocket volume

We also examined the pocket volume changes (PDB 6WQF: Figure S6A,C,E and PDB 6YB7: Figure S10A,C,E). For most states, the pocket volume varies between 200 and 600 ^Å3^. Slight increases in pocket volume are observed for states HE41-HD164 and HE41-HP164 in comparison to other systems. However, it is not clear what pocket volume is optimal for substrate binding. A comparison between the recently published room temperature crystal structure of M^pro11^ and the N3-bound crystal structure indicate regions that undergo significant conformational changes upon ligand binding. Of note, these regions include residues near the P2 site (residues 49-50), the P3-P4 site (residues 166-170), and the P5 loop (residues 190-194) and imply an induced-fit type of ligand-protein interaction. In accord with these previous results, our RMSD/RMSF-per-residue results (Figure S5) reveal a high degree of flexibility/plasticity in these same regions, suggesting that the ligand site volume fluctuations we observed are a signature of active site flexibility.

#### Simulation of the apo state starting from PDB 6YB7

In addition to the room temperature M^pro^ structure,^11^ a low temperature (100 K) crystal structure has been released.^48^ The distinguishing feature in the vicinity of the active site is the altered rotational state of His41. Given the critical functional role His41 plays in the enzymatic activity of M^pro^, we have explored possible differences in the structural stability of M^pro^ upon protonation state variations (Table 1) using 6YB7 as the starting structure. In general, the behaviors observed in the simulations based on PDB entries 6WQF and 6YB7 are strikingly similar, including the behavior of the His164 side chain. The protonation state with low RMSD for both the protein and the catalytic site residues is HD41-HE164 (Figure S8). However, there are several differences; for example, the states CD145-HP41-HD164 and HE41-HE164 appeared to be less structurally stable based on the RMSD in the simulations starting from the 6YB7 structure and in contrast to the simulations based on 6WQF, the His164 in HE41-HP163-HD164 rotates and forms a hydrogen bond with the backbone of Met162. The RMSFs (Figures S5 and S9 for the 6WQF and 6YB7 simulations, respectively), hydrogen bonding (Figures S7 and S11), and water occupancy (Table S3) are mostly the same. There is a reduction in the water occupancy for the HD41-HE164 system in the 6YB7 system; however, the standard deviation is quite large (Table S3).

Collectively, based on RMSD, RMSF, and hydrogen-bonding occupancies in the aforementioned simulations, the most likely apo M^pro^ states appear to be HD41-HE164 and HE41-HE164. However, it should be noted that the differences between states in Group 2 are small, and, therefore, other protonation states of His41-His164 are also possible.

#### Extended simulations

To provide further differentiation between the neutral HD and HE states for His41 and His164, we performed five 250-ns simulations of states HD41-HE164 and HE41-HE164, starting from the room-temperature 6WQF structure. The RMSD of each protein shows that the overall structure is highly structurally stable over the course of the simulations for both states (Figure 6). The RMSD increases slightly from the shorter simulations; however, it remains below 3 ^Å^A. In contrast, significantly larger RMSDs were observed when only the active site was considered, particularly for the state HE41-HE164. Examination of five separate runs shows frequent variations in site RMSD over the course of 250 ns for almost all runs (Figure S12), suggesting that the active site is more flexible than the overall protein structure, as has been seen in previous MD simulations.^27,28^ The RMSF and RMSD per residue (Figure S13) have similar shapes to those from the shorter apo simulations (Figures S5 and S9). Large variations in the pocket volumes were also observed, ranging from 0-800 Å^3^, indicating high site flexibility. Figures S3B,C compare the active sites with small and large volumes and show that the volume is affected by the position of Met165 inside the pocket and by movements of two small loops consisting of residues 47-50 and 188-190 at the pocket opening. These loops correspond to the two largest peaks in the RMSF analysis (Figure S13).

**Figure 6:**
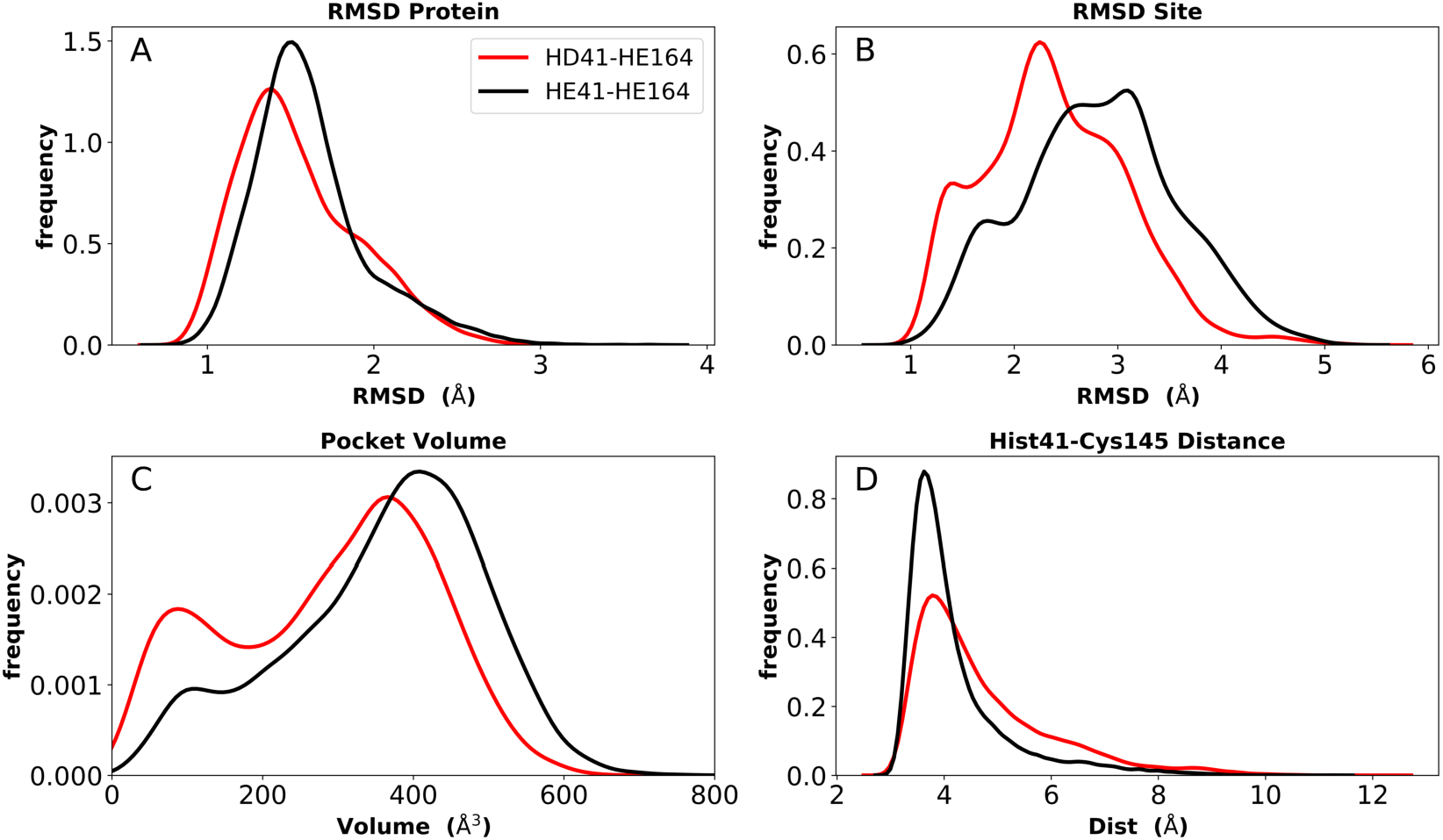
Analysis of longer MD simulations for states HD41-HE164 and HE41-HE164. A) RMSD of protein only. B) RMSD of active site. C) Pocket volume. D) Distance between NE of His41 and S of Cys145.

We monitored hydrogen bonding to residues in the S1 pocket (Figure S14). Overall, the two simulations give similar results, with the HE41-HE164 system having a slightly enhanced occupancy for four of the five interactions. Similar to the shorter apo state trajectories, there is a hydrogen bond between the His164 (donating) and Thr175 (accepting) side chains in the HD41-HE164 state; it is also present in the HE41-HE164 states, but to a much lesser extent, with an occupancy of ~20%. In both systems, the Thr175 hydroxyl side chain interacts with the Asp176 backbone carbonyl, accepting a hydrogen bond. No hydrogen bond between Cys145 and His41 was present in either system, and the distance between NE of His41 and S of Cys145 remained at 4 Å (Figure 6D).

Also analyzed for these longer runs was the interaction of the His41-His164-Asp187 triad with the catalytic water. For HE41-HE164 in all five trajectories for both monomers, and similar to the shorter apo runs, this water was rapidly released into the bulk solvent. Similar to the shorter apo runs, water occupancy in the vicinity of His41, His164, and Asp187 was determined. For the HE41-HE164 system, the site remained occupied by any water molecule for only 21% (±17%) of the trajectory on average, while for HD41-HE164 the occupancy increased to 53% (±28%). Moreover, the largest occupancy average, i.e., the average percentage for a specific water molecule residing in this region, increases from 4% (±4%) for HE41-HE164 to 36% (±30%) for HD41-HE164. Although the standard deviations are large, these results suggest that, relative to HE41-HE164, the HD41-HE164 state has a greater propensity to accommodate a water molecule at this site. Therefore, after taking the longer simulations into account, we propose that the HD-HE164 state is the most likely apo M^pro^ protonation state.

### N3-bound state

The structure of N3, previously developed to inhibit the main protease of multiple coronaviruses, bound to SARS-CoV-2 M^pro^ has been determined (PDB entry 7BQY).^8^ Given that the S1 specificity site for M^pro^ has a nearly absolute requirement for glutamine at position P1 for protein cleavage to occur, ^68^ it is not surprising that the lactam moiety fits well into this site, forming hydrogen bonds with both His163 and Glu166 (Figure 2B). The overall orientation of the ligand is guided by backbone hydrogen bonds between Glu166 and the peptide, as well as the Gln189 side chain. Hydrophobic contacts are also present, as the leucine side chain is inserted into the S2 subsite, a region known to accommodate non-polar substrate groups. ^6^ Overall, these diverse interactions maintain N3 in an orientation competent for the inhibition of M^pro^ enzymatic activity. In light of these stabilizing interactions, we monitored the RMSD of the protein, site, and inhibitor (Figure 7), as well as the RMSF/RMSD per residue, pocket volumes, and hydrogen bonding/hydrophobic contact propensities (Figures S15-S18) across the 12 systems.

**Figure 7:**
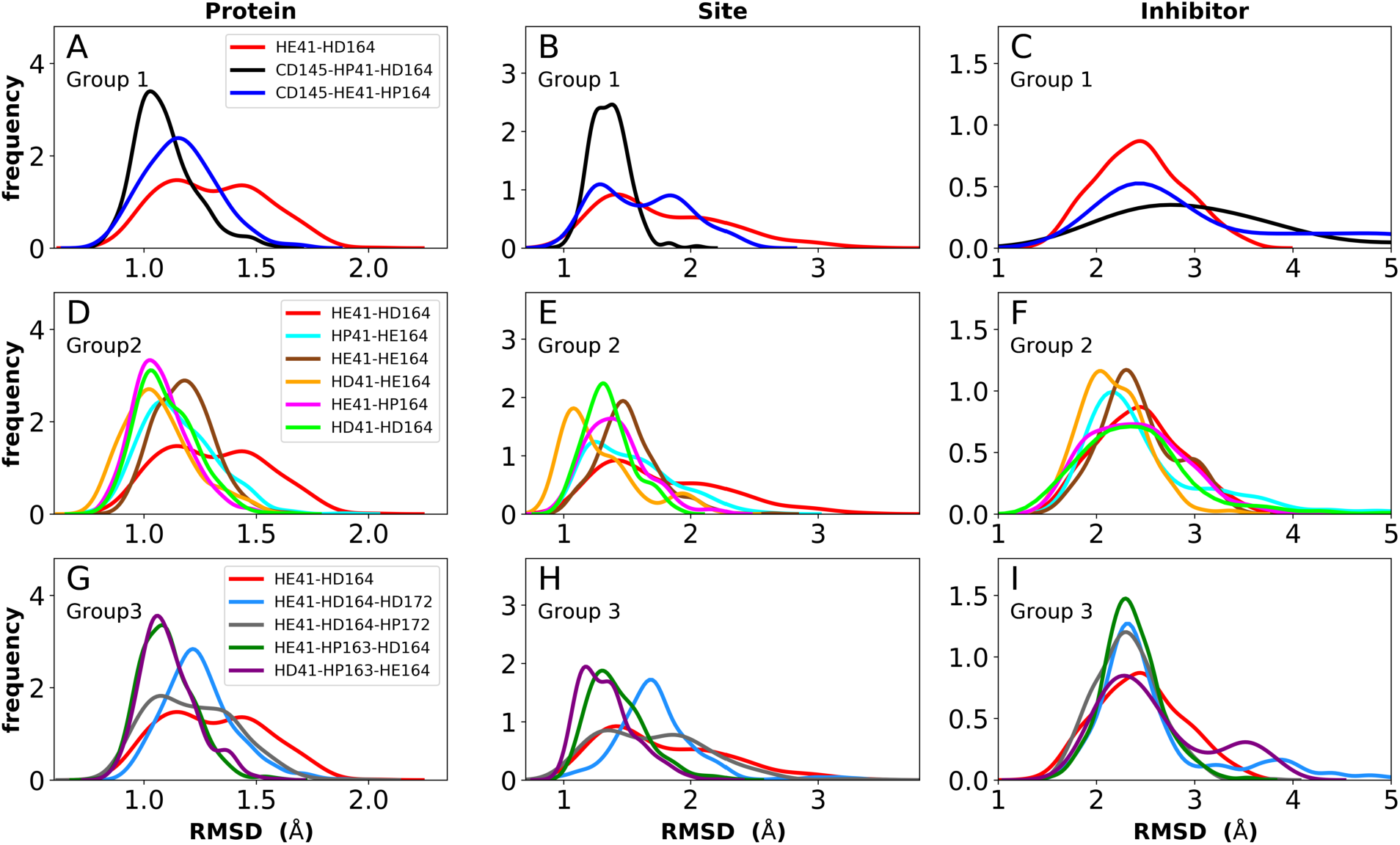
RMSD distributions for the three protonation state groups from simulations of the N3-bound structure (PDB entry 7BQY^8^). A-C) RMSD of (A) protein, (B) active site, and (C) inhibitor for Group 1. D-F) RMSD of (D) protein, (E) active site, and (F) inhibitor for Group 2. G-I) RMSD of (G) protein, (H) active site, and (I) inhibitor for Group 3.

In contrast to the apo systems, RMSD measurements of the inhibitor-bound systems are more sensitive to changes in the protonation states. All states in Group 1 induce high RMSD of either protein and site or the inhibitor, while out of the Group 3 states, only HE41-HP163-HD164 has a low RMSD for these quantities (Figure 7). In addition, for Group 1 states, the distance between Cys145 S and His41 NE was significantly decreased for the state CD145-HP41-HD164 and significantly increased for the state HE41-HD164. (Figure S16B,D,F). This trend was not observed for apo systems. Slightly increased pocket volumes were observed across all systems relative to the apo states (Figure S16A,C,E). In the N3-bound simulations, the total hydrogen bonding interactions and hydrophobic contacts between the ligand and protein were very similar across all the trajectories (Figure S17). The behavior of the hydrogen bonding of the S1 subsite residues is similar to that in the apo simulations in which protonation of His163 ruptures the interaction with Tyr161 (Figure S18). In line with the apo results, the His172-Glu166/His164-Thr175 hydrogen bond is lost in the HD172/HD164 states, respectively, with the Thr175 side chain donating a hydrogen bond to the backbone of Asp176.

Multiple protonation states from Group 2 appear to be feasible (Figure 7) although increased RMSDs were observed for HP41-HE164 and HE41-HE164, and a small decrease in N3-protein hydrogen bonding was observed for HD41-HD164 (Figure S17). Therefore, we conclude that the remaining states in Group 2, namely HD41-HE164, HE41-HP164 and HD41-HD164, are the most structurally stable N3-bound states. Given that HD41-HE164 has a lower RMSD (Figure 7E) for the site residues and the inhibitor (Figure 7F) than either HD41-HD164 or HE41-HP164, and also has robust hydrogen bonding (Figure 2B), and hydrophobic contacts, we propose that this state is the most favorable of the three.

### Ketoamide-bound state

The ketoamide-bound M^pro^ structure (PDB entry 6Y2G) was also simulated in the 12 protonation states listed in Table 1. In contrast to the peptidomimetic N3, the ketoamide carbonyl can accept a hydrogen bond from His41. As illustrated in Figure 2C, this bond occurs only when the NE nitrogen carries a proton. Thus, the ketoamide-bound systems display increased sensitivity to the protonation state of His41 compared to the N3-bound systems. The RMSDs of the protein, site, and inhibitor are illustrated in Figure 8, with RMSF, hydrogen bonding, and hydrophobic contact propensities across the 12 systems also reported (Figures S19-S22).

**Figure 8:**
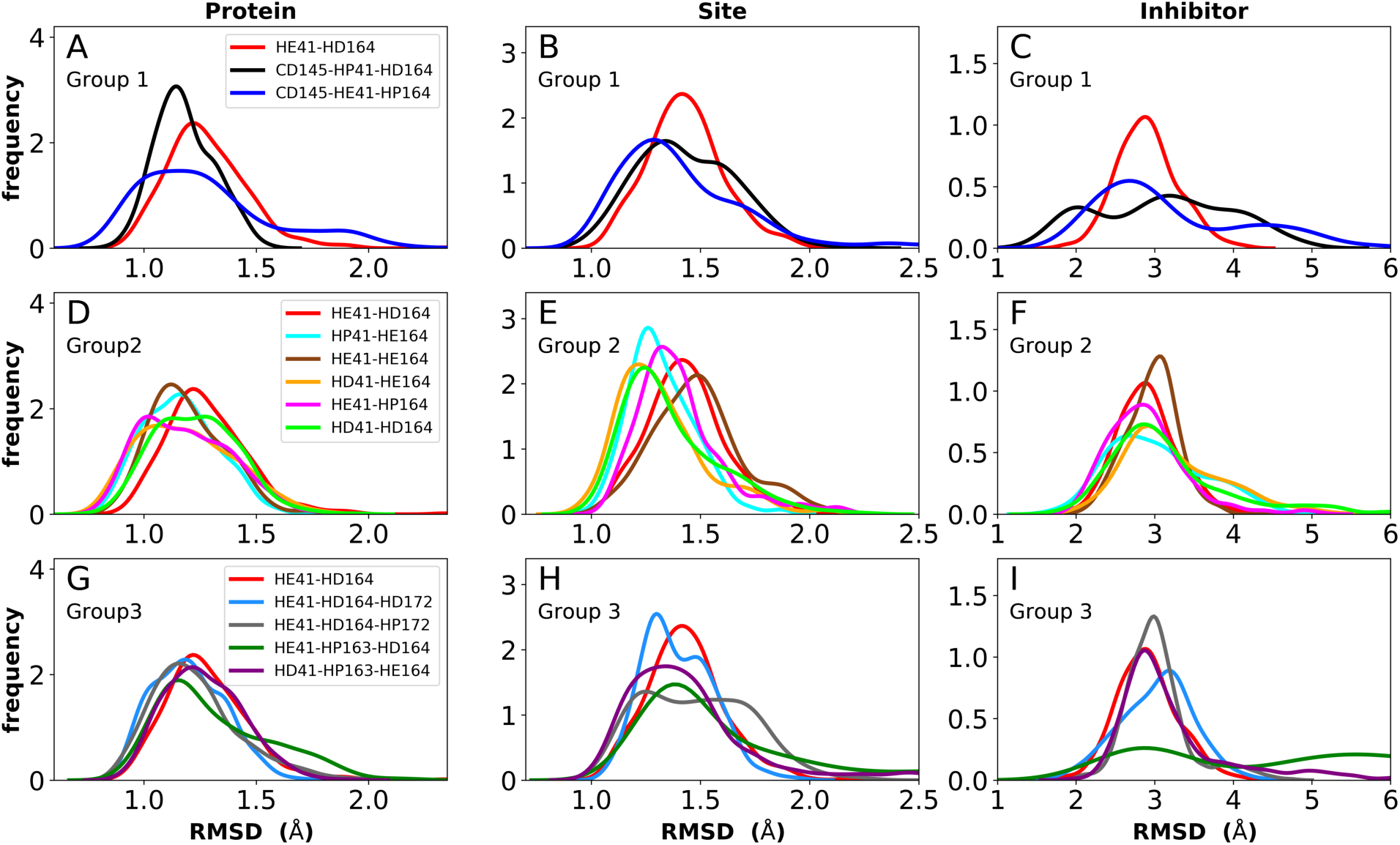
RMSD distributions for the three protonation state groups from simulations of the ketoamide-bound structure (PDB entry 6Y2G). A-C) RMSD of (A) protein, (B) active site, and (C) inhibitor for Group 1. D-F) RMSD of (D) protein, (E) active site, and (F) inhibitor for Group 2. G-I) RMSD of (G) protein, (H) active site, and (I) inhibitor for Group 3.

While the RMSF (Figure S19) profiles and hydrogen bonding of the S1 specificity pocket residues (Figure S22) are similar to the apo and N3-bound simulations, notable differences appear in the RMSD and hydrogen bonding distributions. For Group 1, both states with a deprotonated Cys145 exhibit relatively large and shifted RMSD distributions for the inhibitor (Figure 8C), while the HD41 and HP41 states in Group 2 all display a wide distribution with a slight shoulder at larger values (Figure 8F). These shifts indicate some level of instability of the ligand in the binding pocket. Lastly, of the Group 3 states, in addition to HE41-HD164 only HD41-HP163-HE164 displays an unshifted site and inhibitor RMSD (Figures 8H,I). Of note, for the HD41-HP163-HE164 system, there is a modest reduction in the hydrogen bond count between the ligand and protein (Figure S21). Thus, we conclude that the ketoamide ligand requires a neutral Cys145 and protonation on the NE nitrogen of His41, leaving HE41-HE164, HE41-HD164, and HE41-HP164 as possible protonation states for the ketoamide-bound M^pro^. In line with the apo and N3 simulations, a hydrogen bond between the His164 (donating) and Thr175 side chains is present only in the HE164 and HP164 states, while the His164 side chain frequently rotates in the HD164 states to form a hydrogen bond with the backbone of Met162. This rotation is analogous to that observed in the HD164 apo simulations (Figure 5). Here, we observe that the hydrogen bond to the ketoamide is possible in either conformation of the HE41-HD164 state (Figure 9). In contrast to the other HD164 states, His164 in HE41-HD164 and in HE41-HP163-HD164 also accepts a hydrogen bond from Thr175 (~20% of the time).

**Figure 9:**
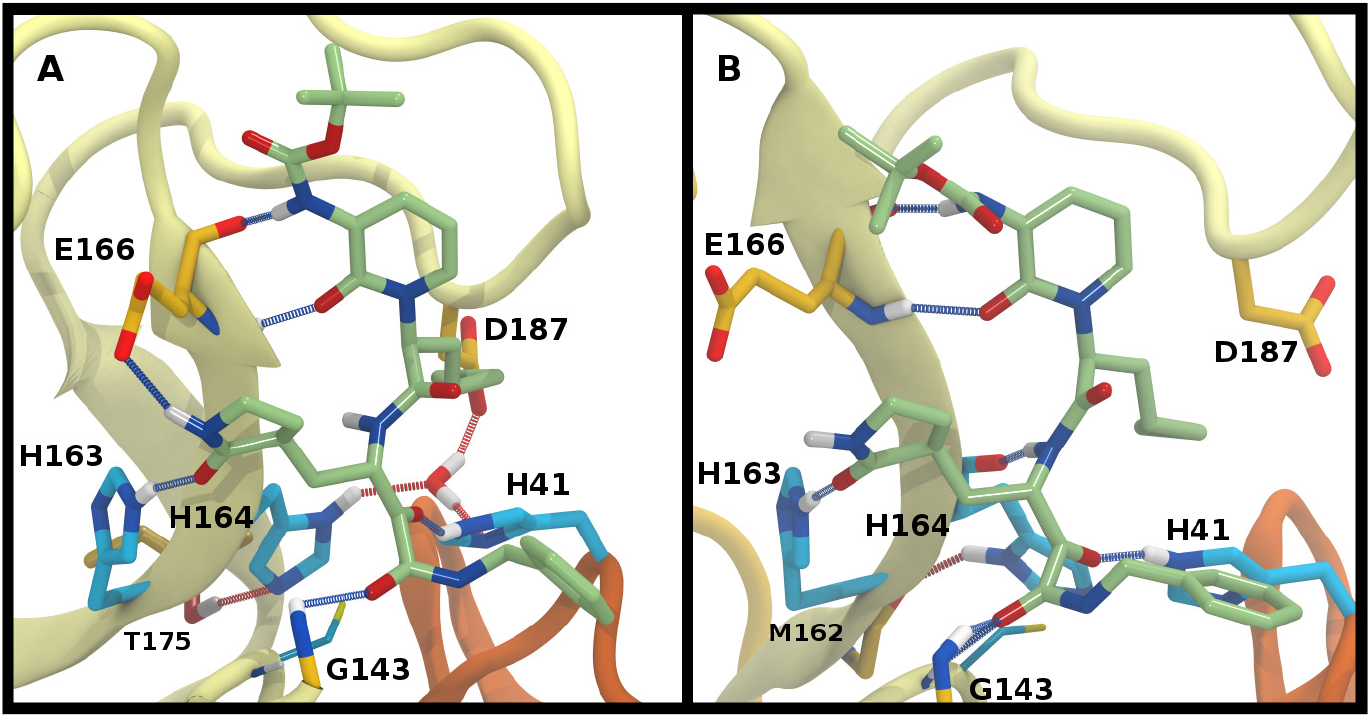
Ketoamide hydrogen bonding in the HE41-HD164 protonation state. In both panels, hydrogen bonds between the ligand (light green licorice) and the protein are indicated with a blue line, while those with water or between protein residues are red. A) Region around the crystallographic water. B) conformation in which the His164 has rotated, making a hydrogen bond with the backbone of Met162. The crystallographic water has been released.

Notably, protonation states that have low RMSD in N3-bound and apo structures, e.g., HD41-HE164 and HE41-HP163-HD164, showed increased displacement of the ketoamide inhibitor, as measured by its RMSD (Figure 8) and significantly reduced total hydrogen bonding to M^pro^ (Figure S21). In particular, the most dramatic effect observed is for the HE41-HP163-HD164 system. Although the HD41-HE164 state still has low RMSD for both the protein and the active site, increased RMSD of inhibitor and decreased inhibitor-protein hydrogen bonding indicates that this protonation state is structurally unstable for ketoamide-bound M^pro^. This observation is in contrast to apo and N3-bound simulations, where the HE41-HD164 state was not particularly stable structurally instead, and appeared to be destabilizing for the N3-bound state due to high RMSD. This is most likely a result of the sensitivity of the ketoamide-bound simulations to the presence of the HE41 carbonyl hydrogen bond, not present in the N3-bound structure (Figure 2B,C).

### Free-energy calculations for His41 and His163 protonation states

As observed in the preceding discussion of possible protonation states, the state of His41 is quite sensitive to the presence of the ligand, favoring HD41 in the N3-bound system and HE41 in the ketoamide system. In order to further delineate the relative stability between HD41 and HE41 in the two systems, free-energy perturbation calculations (FEP) were performed (Table 2). In addition, we investigated the change in free energy for HE163→HP163. The FEP calcuations were only used for the two histidines that directly interact with the bound ligands. Our results confirmed that while HE41 is preferred by ketoamide by 1.3 kcal/mol, the HD41 state is preferred by N3 by 0.8 kcal/mol (both numbers per monomer). Although small (1-2 kT), the energetic differences for the two states are statistically significant (Table 2). In addition, the difference in structural stability for the two states is also supported by changes in inhbitor RMSD and hydrogen bonding with the protein.

**Table 2:** The relative protein-ligand binding free energy, ΔΔG°, corresponds to the difference in binding affinities between the initial and the target (WT) enzyme, i.e., 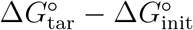, which is equal to the difference in alchemical free energies between the bound and unbound states, i.e., 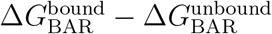 (see Table S4). The ΔΔ*G*° values reflect the concomitant amino-acid protonation state change in both monomers of the homodimeric enzyme. The values supplied in parentheses represent the relative protein-ligand binding free energy per monomer. All energies are in kcal/mol.

| transformation | substrate | $\Delta\Delta G^\circ$ |
| --- | --- | --- |
| HD41→HE41 | ketoamide | $-2.7 \pm 0.3$ ($-1.3 \pm 0.3$) |
| HE163→HP163 | ketoamide | $+6.7 \pm 1.4$ ($+3.4 \pm 1.4$) |
| HD41→HE41 | N3 | $+1.5 \pm 0.1$ ($+0.8 \pm 0.1$) |
| HE163→HP163 | N3 | $+3.9 \pm 1.0$ ($+1.9 \pm 1.0$) |

Although the HE163→HP163 transformation was unfavorable for both inhibitors (Table 2), the magnitude of free energy change was larger for ketoamide by 1.5 kcal/mol. This is in qualitative agreement with our analysis of hydrogen bonding for the two compounds in HP163 states, which shows a larger loss in hydrogen bonding for ketoamide (~2.5 hydrogen bonds lost on average) than in the corresponding N3 state (~0.5 hydrogen bonds lost; Figures S17 and S21) relative to the HE41-HD164 state.

## Conclusions

To combat COVID-19, there is tremendous effort aimed at developing both vaccines and antiviral drugs against its causative agent, SARS-CoV-2. The main protease of SARS-CoV-2 (M^pro^), a homodimeric cysteine protease, has emerged as an attractive target for drug design given 1) its critical role in early stages of the virus replication cycle, 2) its similarity to main proteases from other betacoronaviruses, thereby leveraging earlier antiviral development ef-forts, and 3) its unique substrate specificity, cleaving primarily after a glutamine, a target unknown for other host cell proteases.^5–7^

A number of crystal structures of M^pro^ have been determined, including the apo state as well as bound to covalent, non-covalent, and fragment-based inhibitors.^7,8,11^ The substrate binding site in M^pro^ consists of a catalytic Cys145-His41 dyad, as well as several histidine residues in close proximity to the catalytic site including His172, His163, and His164. These residues, along with Glu166 and the N-terminus of the other monomer, Ser1, form an interlocking hydrogen-bonded pocket, which is the target of most designed inhibitors. For computational drug design strategies to be effective, the details of the structure and dynamics of these residues are required, particularly under different protonation states. Here we have enumerated these states and determined the effects of protonating various residues for M^pro^ in both the apo form as well as ligand-bound complexes using MD simulations.

In the present study, we have demonstrated that the combination of protonation states for histidines in or near the catalytic site can have a profound impact on M^pro^’s structural stability, as measured by RMSD and RMSF, as well as the hydrogen bonding occupancies, hydrophobic contacts, and catalytic subsite characteristics, e.g., available pocket volume. Examining these properties, we conclude that the protonated His41/deprotonated Cys145 state of the catalytic dyad is structurally unstable in the crystal structure conformations as exemplified by increased RMSD, altered hydrogen bonding patterns, and unbinding of the inhibitors. However, this state may exist as a transient reaction intermediate.

We have also shown that His163 and His172 protonation states other than HE result in significant perturbations to several hydrogen bonds compared to crystal structure conformations. Although HE protonation was expected for these two residues based on structural data, concurrent HD protonation could not be excluded without the results from our MD simulations, particularly for His172, which is thought to be protonated in M^pro^ of SARS-CoV.^17^ In addition, decreased pocket volume was frequently observed in the simulations for the HP163 state; free-energy calculations showed decreased affinity for both inhibitors in this state, which qualitatively tracks the hydrogen bonding reductions observed. The sensitivity of the M^pro^ structure to the protonation states of His163 and His172 may explain why M^pro^ is inactive at low pH.^18,21^ At low pH, HP163 is expected to interfere with substrate binding due to decreased pocket volume and substrate affinity, whereas fluctuation of the N-finger and decreased hydrogen bonding within the protein, induced by the HP172 state, could interfere with catalysis.

In contrast to the other histidines, we determined that multiple protonation combinations are feasible for the His41-His164 pair. It is possible that these residues adopt different protonation states during protein cleavage. We conclude that HD41-HE164 appears to be the more stable state for the apo and N3-bound structures, whereas HE41-HD164 is more structurally stable for the ketoamide-bound structure. This change in protonation state preference of M^pro^ for the two inhibitors was also confirmed with free-energy calculations (Table 2). Unlike N3, ketoamide has two carbonyl groups around its reactive site, facilitating additional bonding to M^pro^ (Figure 2C). In fact, the structural data indicate that protonation of the NE nitrogen in His41 is preferred in order to provide an interaction with the ketoamide carbonyl. Consequently, ketoamide has slightly better in vitro inhibition activity than N3 (5 *μM* vs. 17 μM),^7,8^ although different reaction mechanisms of covalent bond formation with Cys145 for Michael acceptors and ketoamides as well as their ability to go through membranes can also contribute to differences in activity. Nevertheless, additional optimization to improve their potency is desirable for both compounds. The different protonation states of M^pro^ histidines will need to be considered for such optimization efforts, as well as for the rational design of other inhibitors. We expect that other peptidomimetic Michael acceptors and *α*-ketoamides will have the same protonation state preferences as N3 and ketoamide, respectively. However, in silico high-throughput screens of novel potential inhibitors will benefit from being performed on each of the protonation states considered here to be feasible in the presence of a ligand.

## Acknowledgement

JCG acknowledges support from the National Institutes of Health (R01-AI148740). JMP was supported by the DOE Office of Science through the National Virtual Biotechnology Laboratory, a consortium of DOE national laboratories focused on response to COVID-19, with funding provided by the Coronavirus CARES Act. This research used resources of the Oak Ridge Leadership Computing Facility at the Oak Ridge National Laboratory, which is supported by the Office of Science of the U.S. Department of Energy under Contract No. DE-AC05-00OR22725. This work also used resources, services, and support provided via the COVID-19 HPC Consortium (https://covid19-hpc-consortium.org/), which is a unique private-public effort to bring together government, industry, and academic leaders who are volunteering free compute time and resources in support of COVID-19 research. Additional resources were provided by the Partnership for an Advanced Computing Environment (PACE) at the Georgia Institute of Technology. This research used resources at the Spallation Neutron Source and the High Flux Isotope Reactor, which are DOE Office of Science User Facilities operated by the Oak Ridge National Laboratory.

## Supporting Information Available

Extended analysis of all systems and additional details on force-field parametrization.

