## Supplemental Information for "Inhibitor binding influences the protonation states of histidines in SARS-CoV-2 main protease"

### Parametrization of ketoamide

The initial parameters were taken from CGenFF website, which also provides penalties for assigned partial charges and parameters. At first, only charges with penalties higher than 10 were optimized by matching the energies and distances of interactions with water molecules at QM and MM levels, as advised for CHARMM compatible parameters. A few atoms with penalties less than 10 were added for the final charge optimization in order to improve the quality of the fit. The force field Toolkit (ffTK) was used for all charge and parameter optimizations and Gaussian16 was used for all QM calculations. Because the ketoamide molecule was too large for calculations at the MP2 level of theory, we used a smaller model (ketoamide model I) that still contained all atoms needing optimization (Figure S1). All geometries were optimized at MP2/6-31G* level of theory and same level of theory was used for frequency calculations and dihedral scans. Water interactions were optimized at the HF/6-31G* level of theory, in accordance with CGenFF guidelines for these calculations. The HF energies were scaled by a factor of 1.16 as recommended for neutral compounds. For most atoms, a reasonable fit (error < 0.7 kcal/mol) was achieved (Table S1). It should be noted that atoms with larger errors in QM interaction energies, namely C24, N15 and C40 (Figure S2), do not interact with the protein.

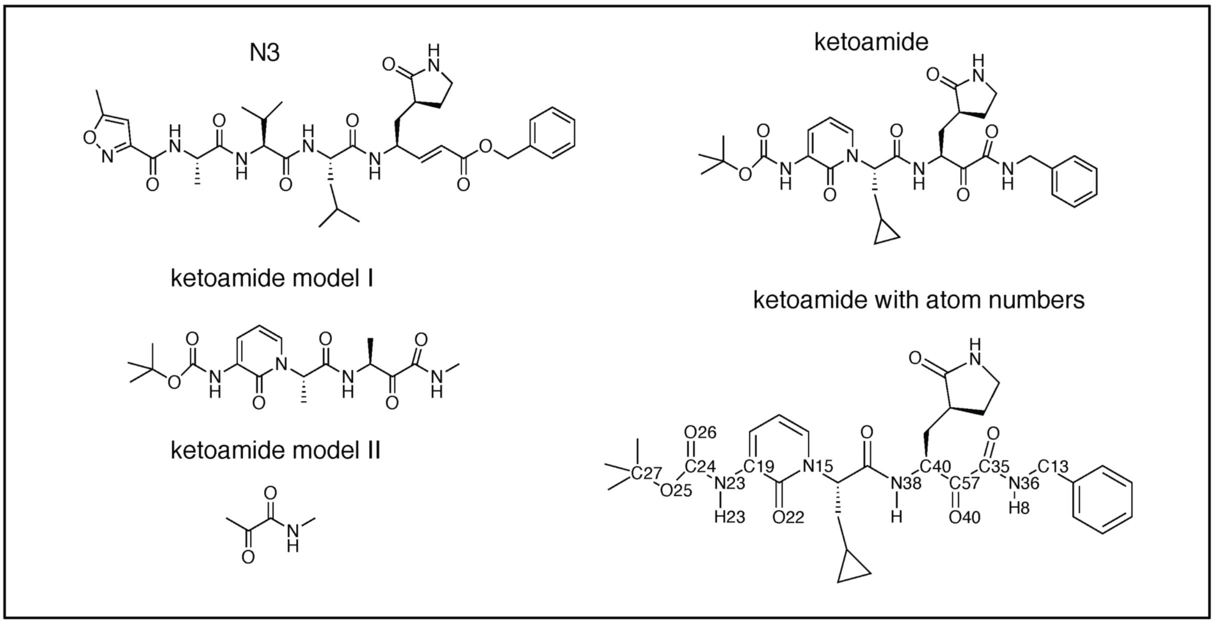


Figure S1: Chemical structures of N3, ketoamide and ketoamide models used for charge optimization. Names of atoms with optimized charges are also shown.

Table S1: List of atom names, QM and MM interaction energies, error in interaction energies and the fitted charges for all atoms with optimized charges. All energies are in kcal/mol and QM energies are scaled by 1.16.

| Atom name | QM energy (scaled) | MM energy | Error | Charge |
|---|---|---|---|---|
| C24 | -1.057 | 0.135 | 1.192 | 0.338 |
| O26 | -5.569 | -5.809 | -0.239 | -0.451 |
| N23 | -0.722 | -0.618 | 0.104 | -0.255 |
| C19 | -0.802 | -0.861 | -0.059 | 0.26 |
| N15 | -0.191 | -2.783 | -2.592 | -0.333 |
| N38 | -1.681 | -2.287 | -0.606 | -0.476 |
| C40 | 1.07 | 0.033 | 1.103 | 0.078 |
| C57 | -3.101 | -2.6 | 0.501 | 0.515 |
| O40 | -5.818 | -5.607 | 0.211 | -0.364 |
| C35 | -1.738 | -1.492 | 0.247 | 0.243 |
| N36 | -0.648 | -1.016 | -0.367 | -0.282 |
| C13 | -2.603 | -1.925 | 0.678 | -0.082 |
| H8 | -5.932 | -5.989 | -0.057 | 0.284 |
| H23 | -0.755 | -0.474 | 0.281 | 0.261 |
| O22 | -6.799 | -6.434 | 0.365 | -0.444 |
| O25 | -3.502 | -3.179 | 0.323 | -0.327 |
| C27 | -1.869 | -1.182 | 0.687 | 0.256 |

Bond, angles and dihedral terms were fitted using ffTK. Only the ketoamide part of the molecule needed re-optimization of bond and angle terms. Because frequency calculations are more memory intensive than geometry optimizations, our model of ketoamide needed to be reduced further (ketoamide model II). Only one bond term was fitted, matching the target equilibrium QM value, while all the angle terms were within 0.05° of their target equilibrium QM values. All of the energies for bond and angle displacements were within 0.06 kcal/mol. The final results for optimization of dihedral terms are shown in Fig S2, comparing the QM and MM energies of the dihedral scans. Good agreement is observed between MM and target QM data. All of the fitted parameters are added to the attached parameter file.

### Calculations of the pocket volume

In the Epock software package, the pocket cavity is defined by the user using a number of spheres. The centers of these spheres are based on centers of specified selections, and radius sizes are controlled by the user. The selections and sphere sizes used to define the pocket in

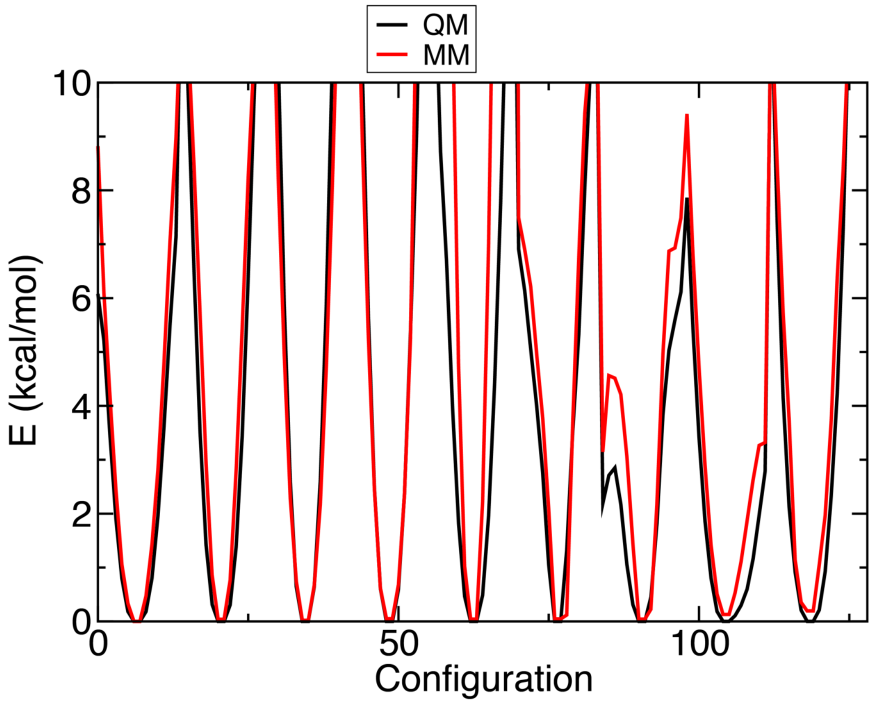


Figure S2: Comparison of QM (black) and MM (red) energies of dihedral scans performed for ketoamide in order to optimize the dihedral parameters.

each monomer are given in Table S2 and Figure S4.

Table S2: Selections used to define the pocket of each monomer for volume calculations in Epock.

| Complete residues | Size (Å) |
|---|---|
| 41 145 | 5.0 |
| 166 168 189 | 4.0 |
| 166 163 | 3.0 |
| 41 49 | 3.0 |
| 49 140 | 5.0 |
| Side chain only | Size (Å) |
| 140 141 189 | 5.0 |
| Backbone only | Size (Å) |
| 49 165 | 4.0 |
| 27 142 143 | 4.0 |

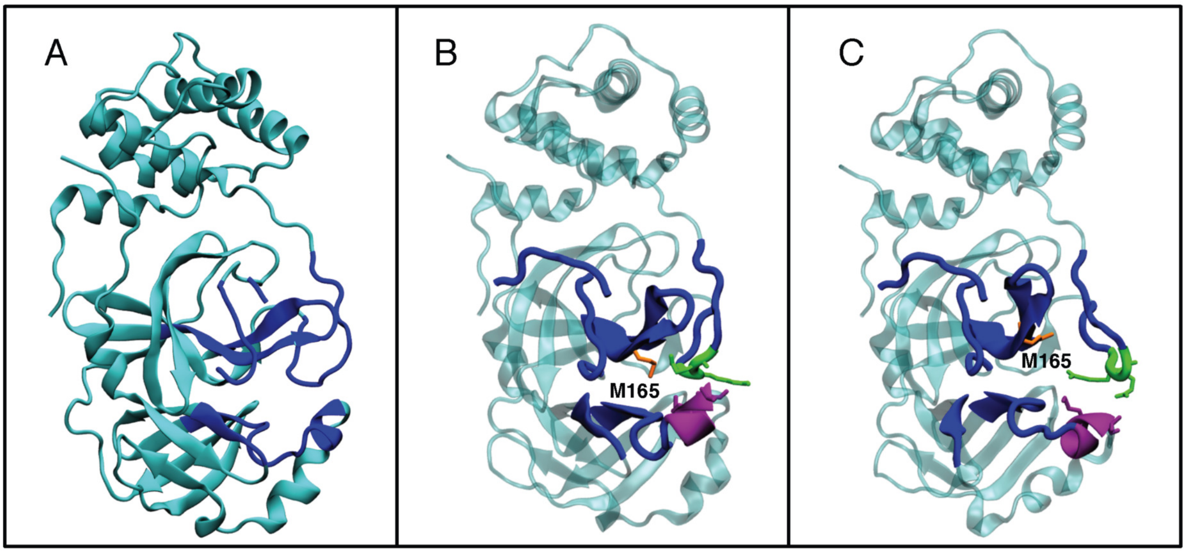


Figure S3: A) Structure of one monomer (PDB code 6WQF) with the defined active site shown in blue. B) Snapshot of a structure with low pocket volume (10 Å) from the extended simulations of HE41-HD164 state. The active site is shown in blue, with exceptions of M165 in orange, and the loops with residues 188-190 and 48-50 are shown in green and purple, respectively. Met165 moves into the center of the pocket, and the purple and green loops have moved close together. C) Snapshot of a structure with high pocket volume (605 Å) from the extended simulations of the HE41-HD164 state. Same colors as in B are used. Met165 is seen to move to the side of the pocket and the green and purple loops move farther apart.

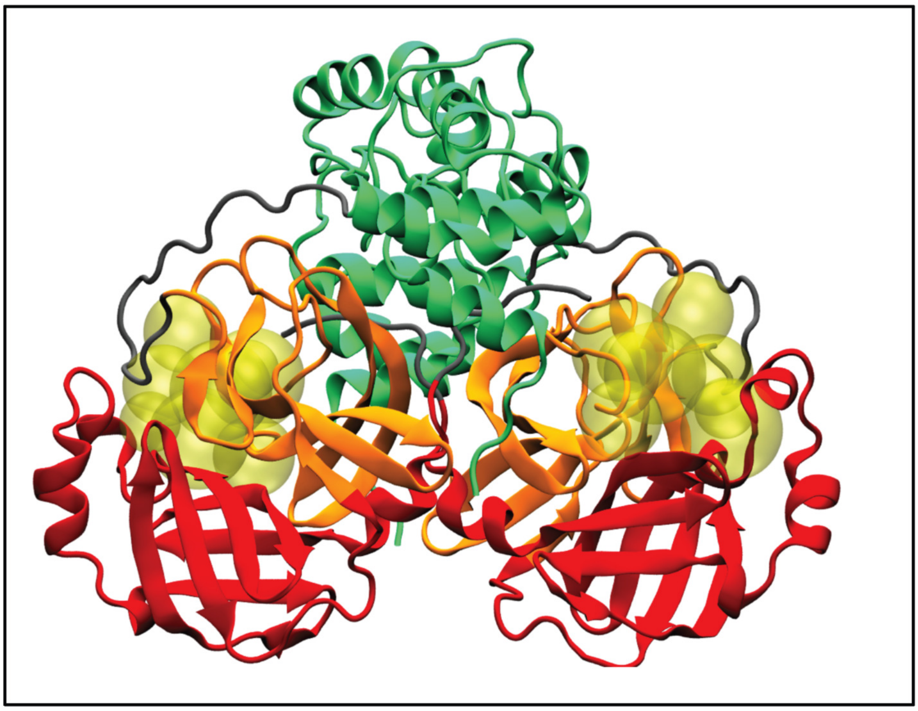

Figure S4: Structure of $M^{pro}$ dimer with domains I, II and III colored in orange, red and green, respectively. The spheres used to define the active site pockets are shown in yellow.

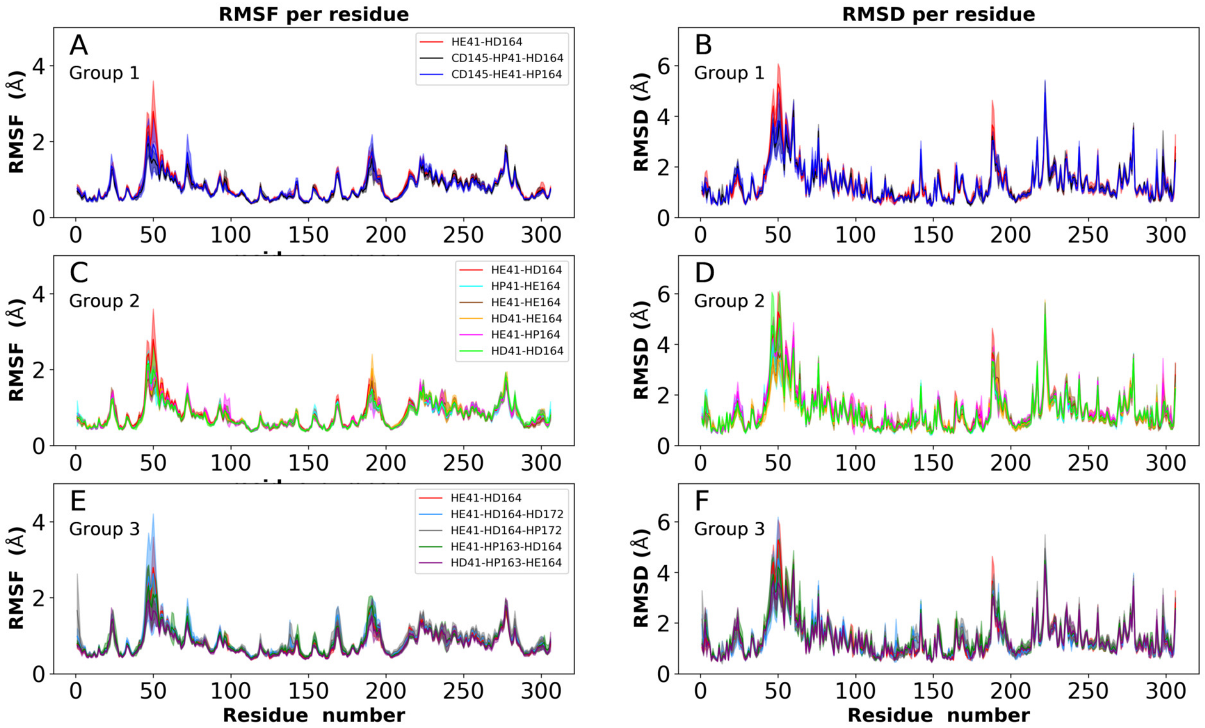


Figure S5: RMSF of $C_\alpha$ atoms (left) and RMSD of whole residues (right) for all simulations of apo structure 6WQF.

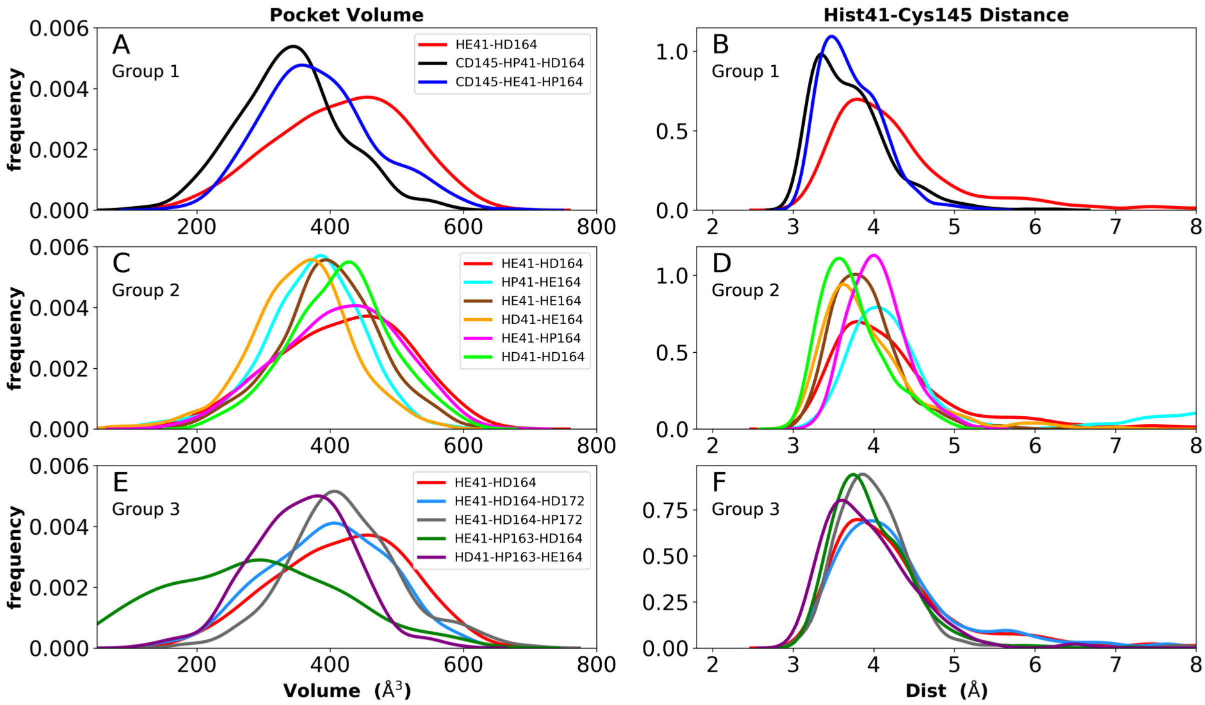


Figure S6: Distributions of pocket volume (left) and distance between NE and S in the catalytic residues His41 and Cys145 (right) for apo structure 6WQF.

Table S3: Percent occupancy of the catalytic water site. Averages based on the last 15 ns of the trajectories with standard deviations in parentheses.

| State (PDB code) | apo(6WQF) | apo(6YB7) |
|---|---|---|
| **Group 1** | | |
| CD145-HE41-HP164 | 37 (±18) | 26 (±4) |
| CD145-HP41-HD164 | 18 (±27) | 4 (±7) |
| HE41-HD164 | 15 (±28) | 19 (±33) |
| **Group 2** | | |
| HE41-HD164 | 15 (±28) | 19 (±33) |
| HD41-HD164 | 76 (±33) | 70 (±38) |
| HD41-HE164 | 80 (±25) | 52 (±50) |
| HE41-HE164 | 14 (±8) | 23 (±23) |
| HE41-HP164 | 34 (±12) | 27 (±7) |
| HP41-HE164 | 30 (±22) | 10 (±15) |
| **Group 3** | | |
| HE41-HD164 | 15 (±28) | 19 (±33) |
| HD41-HP163-HE164 | 32 (±17) | 24 (±25) |
| HE41-HD164-HD172 | 2 (±1) | 0 |
| HE41-HD164-HP172 | 13 (±17) | 34 (±45) |
| HE41-HP163-HD164 | 48 (±43) | 3 (±3) |

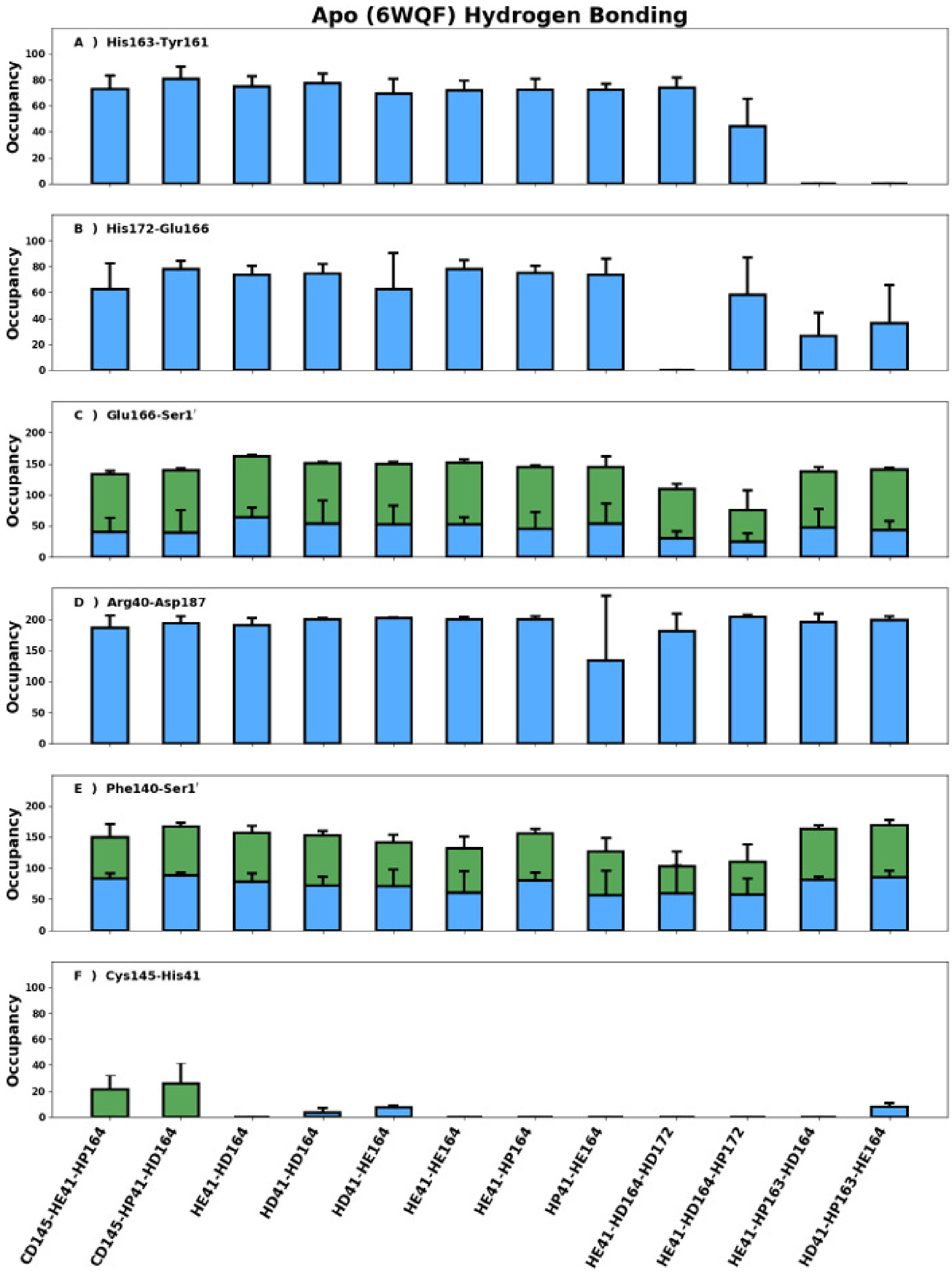


Figure S7: Hydrogen Bonding occupancies and standard deviations in 6WQF based trajectories. A) His163-Tyr161, B) His172-Glu166, C) inter-monomer Glu166-Ser1′ with blue/green indicating the serine side chain/backbone respectively, D) the Arg40-Asp187 salt bridge/charge reinforced hydrogen bond, E) inter-monomer interaction Phe140-Ser1′ with blue/green indicating Ser1′ acting as an acceptor/donor, and F) the catalytic dyad residues with blue/green indicating the sulfur acts as donor/acceptor. In all figures, unless otherwise stated, His163 and His172 are HSE and standard deviations are indicated with bars. Also note, occupancies can be greater than 100% in cases where more than one hydrogen bond can be formed.

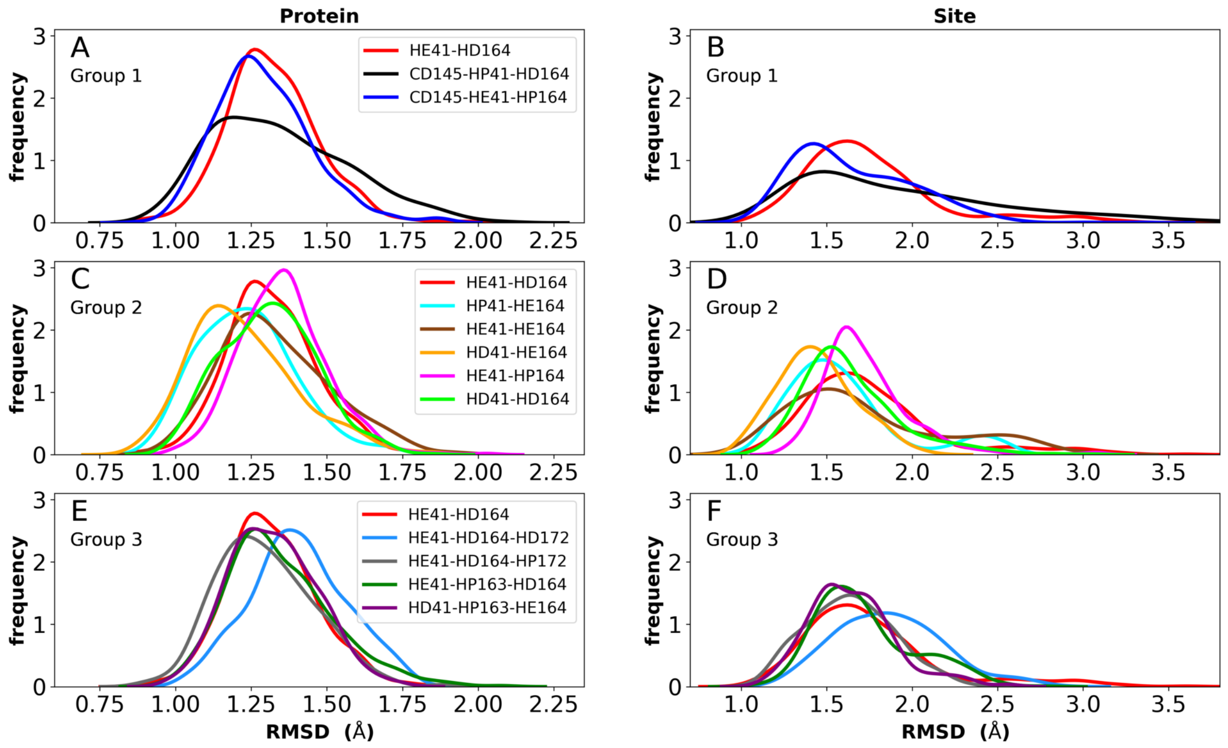


Figure S8: Distribution plots of RMSD for the three protonation state groups from simulations of the apo structure with PDB code 6YB7. A) RMSD of protein for Group 1. B) RMSD of active site for Group 1. C) RMSD of protein for Group 2. D) RMSD of active site for Group 2. E) RMSD of protein for Group 3. F) RMSD of active site for Group 3.

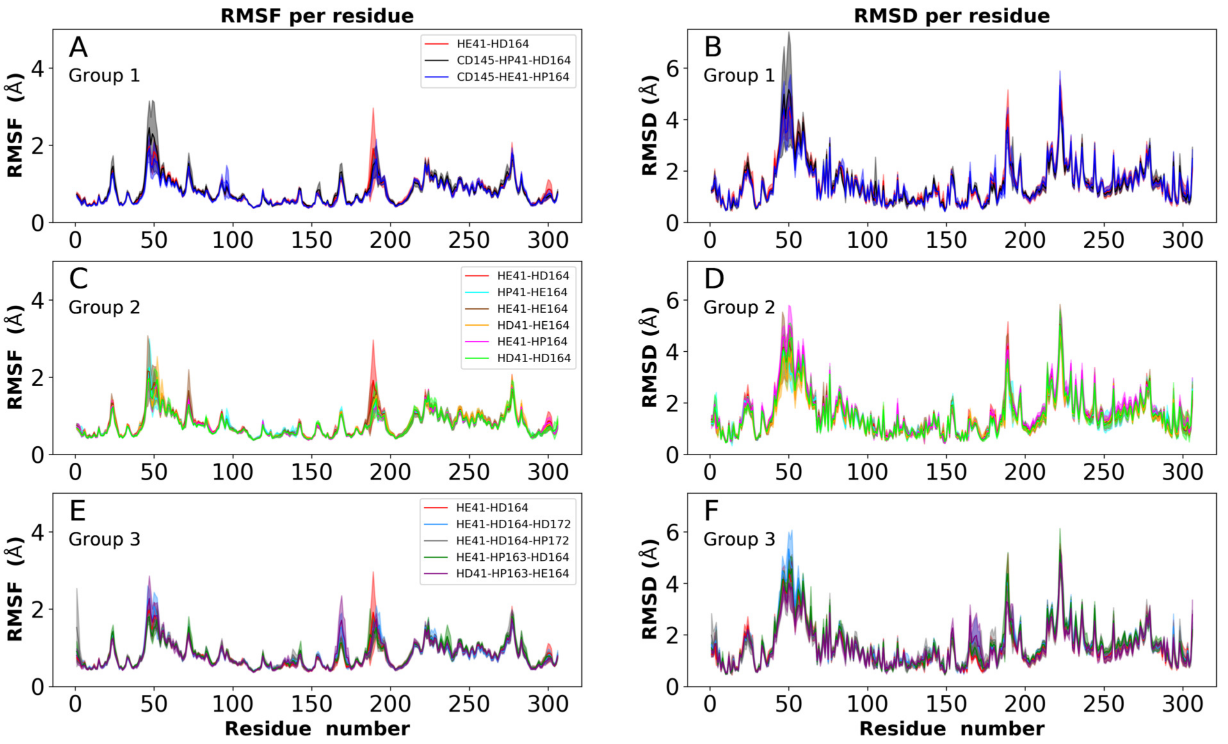


Figure S9: RMSF of $C_\alpha$ atoms (left) and RMSD of whole residues (right) for all simulations of apo structure 6YB7.

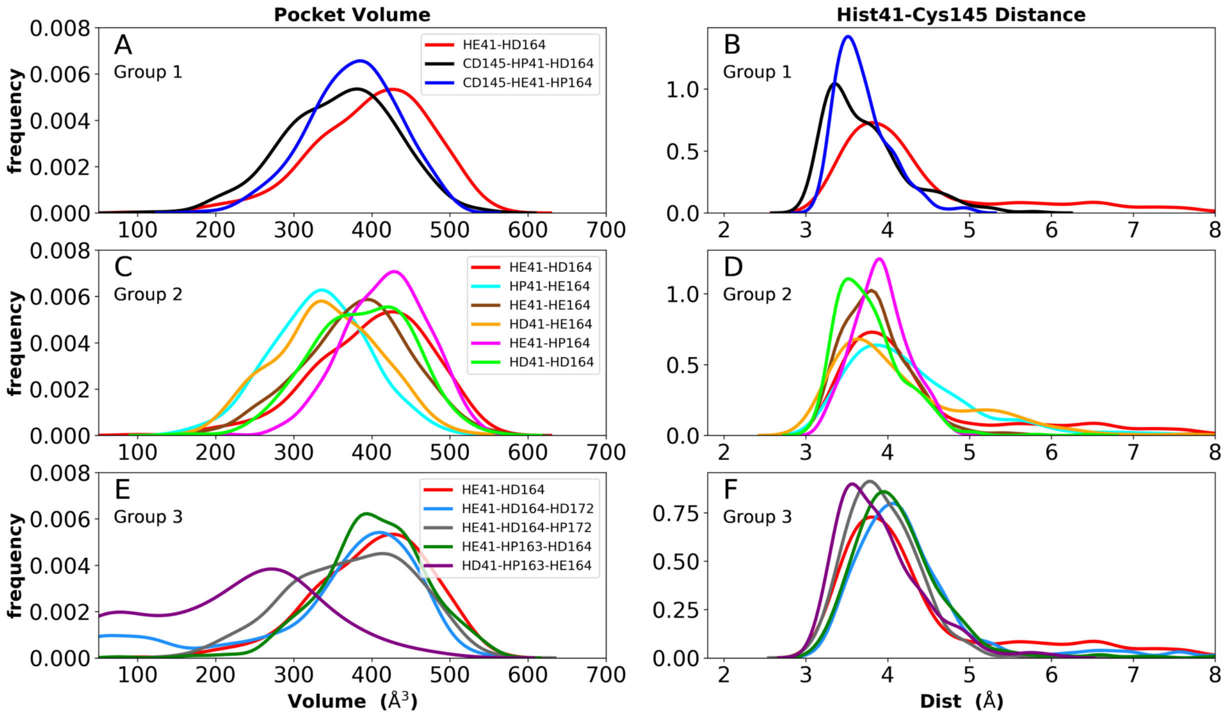


Figure S10: Distributions of pocket volume (left) and distance between NE and S in the catalytic residues His41 and Cys145 (right) for apo structure 6YB7.

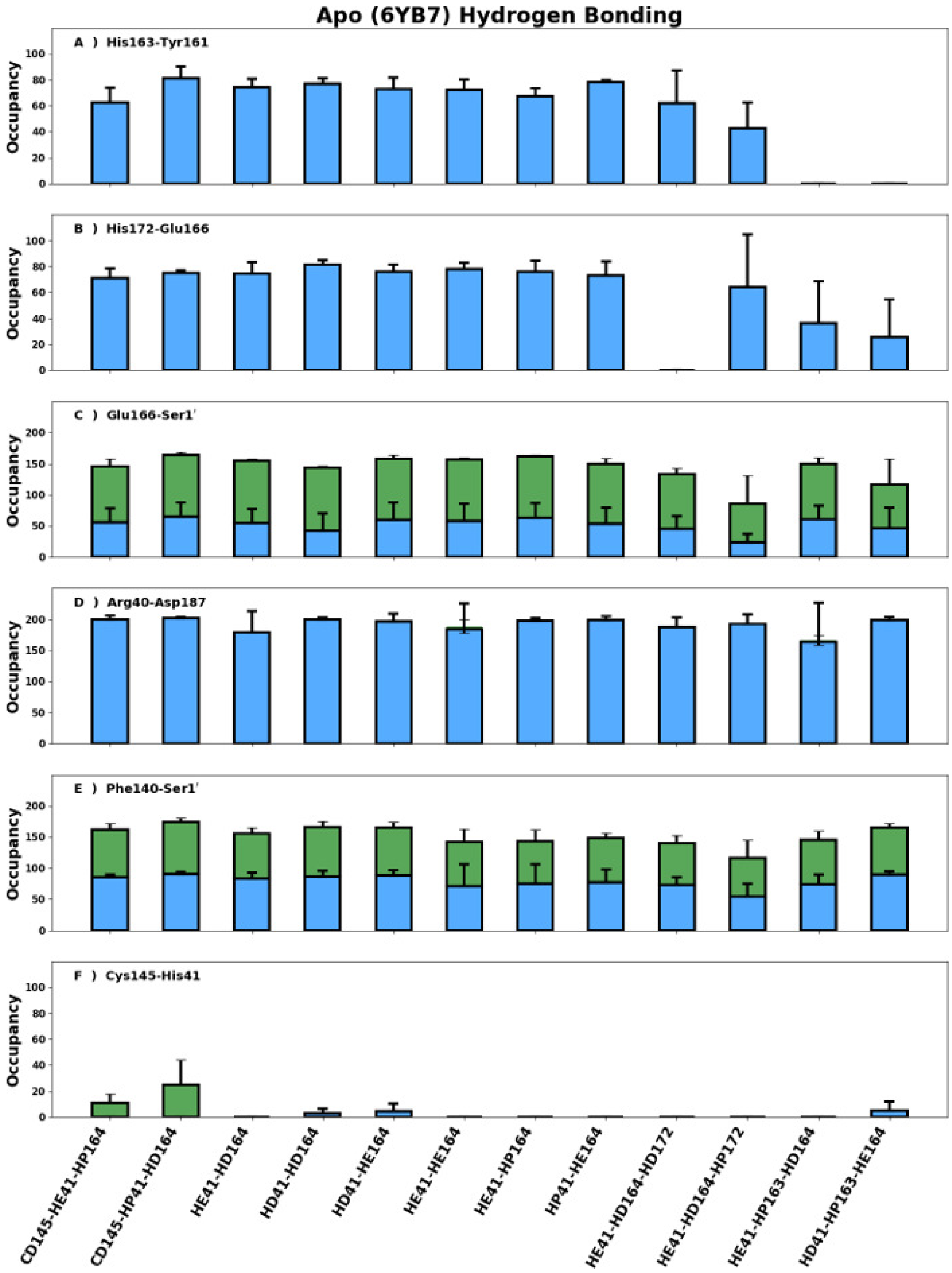


Figure S11: Hydrogen Bonding occupancies and standard deviations in 6YB7-based trajectories. A) His163-Tyr161, B) His172-Glu166, C) inter-monomer Glu166-Ser1′ with blue/green indicating the serine side chain/backbone respectively, D) the Arg40-Asp187 salt bridge/charge reinforced hydrogen bond, E) inter-monomer interaction Phe140-Ser1′ with blue/green indicating Ser1′ acting as an acceptor/donor, and F) the catalytic dyad residues with blue/green indicating the sulfur acts as donor/acceptor. In all figures, unless otherwise stated, His163 and His172 are HSE and standard deviations are indicated with bars. Also note, occupancies can be greater than 100% in cases where more than one hydrogen bond can be formed.

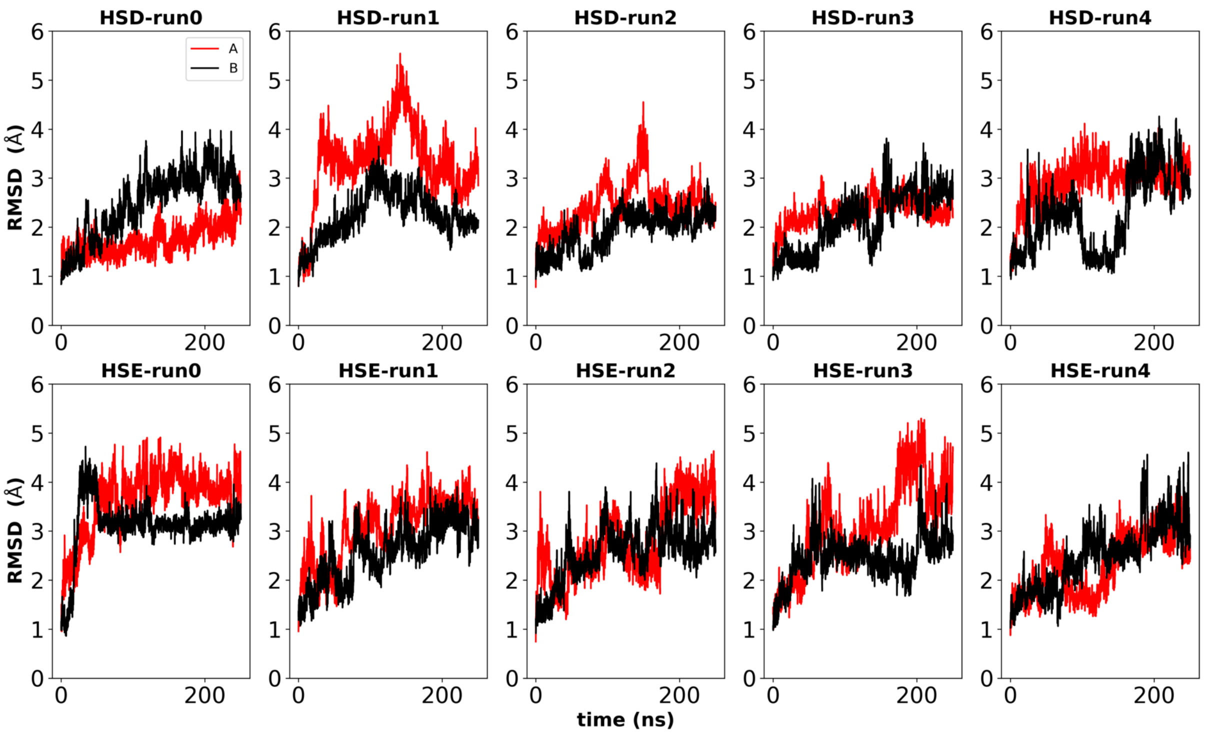


Figure S12: RMSD of the active site for each separate run and each monomer (A and B) for the extended apo simulations.

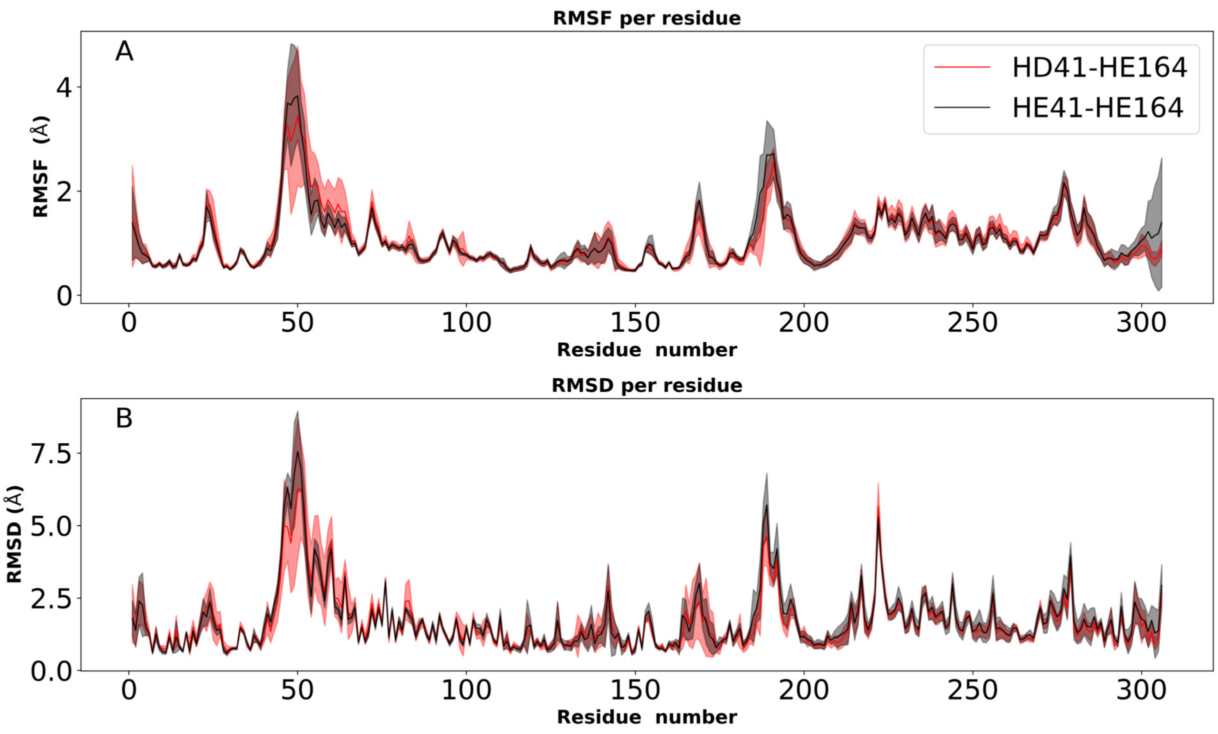


Figure S13: A) RMSF of $C_\alpha$ atoms for the extended simulations starting from apo structure 6WQF. B) RMSD for each residue from the same simulations.

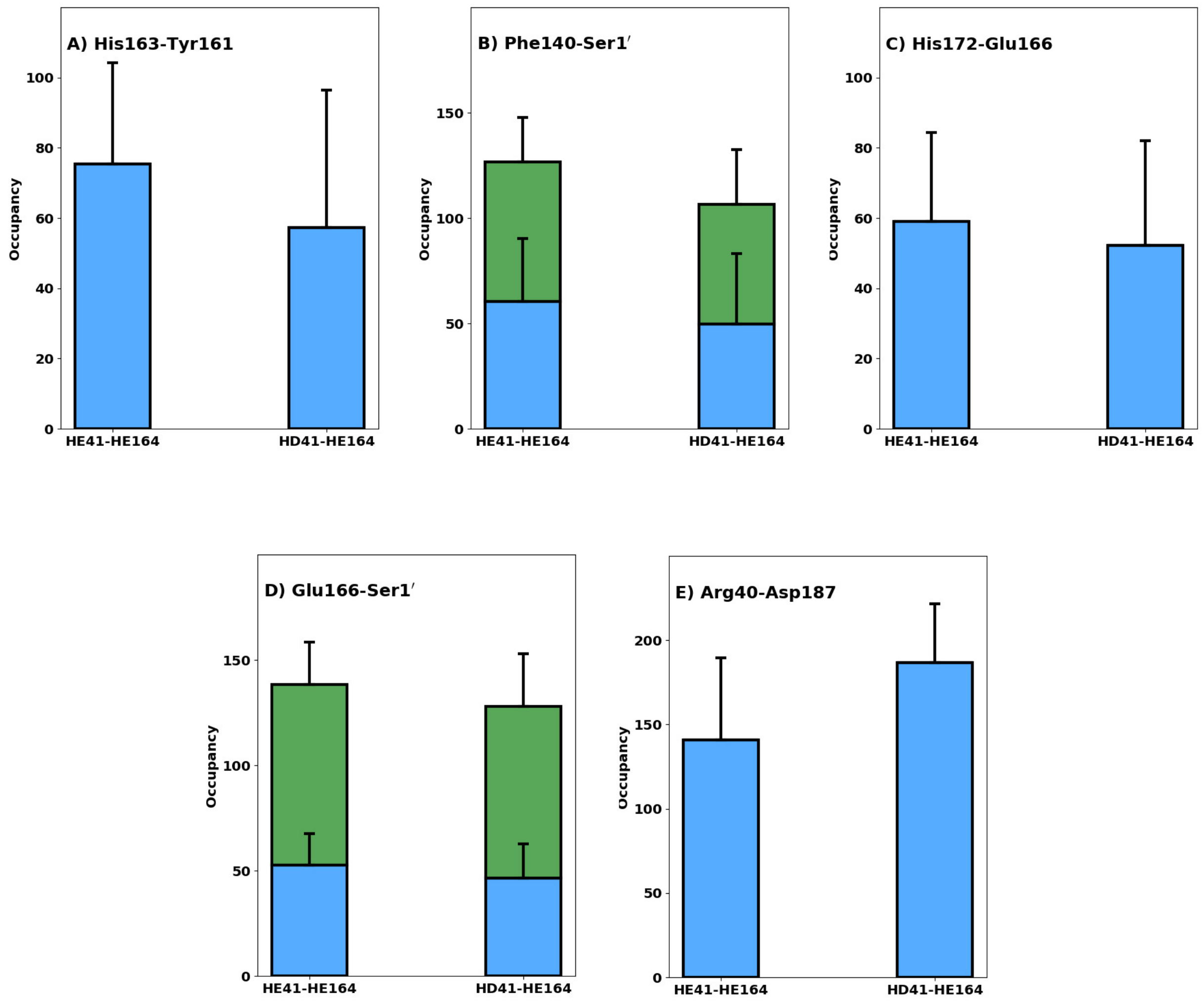


Figure S14: Hydrogen Bonding interactions for the HE41-HE164 and HD41-HE164 systems based on five independent trajectories of 250 ns each. A) His163-Tyr161, B) inter-monomer interaction Phe140-Ser1′ with blue/green indicating serine acting as an acceptor/donor respectively, C) His172-Glu166, D) inter-monomer Glu166-Ser1′ with blue/green indicating the S1 side chain/backbone, and E) the Arg40-Asp187 salt bridge/charge reinforced hydrogen bond. Standard deviation indicated with bars.

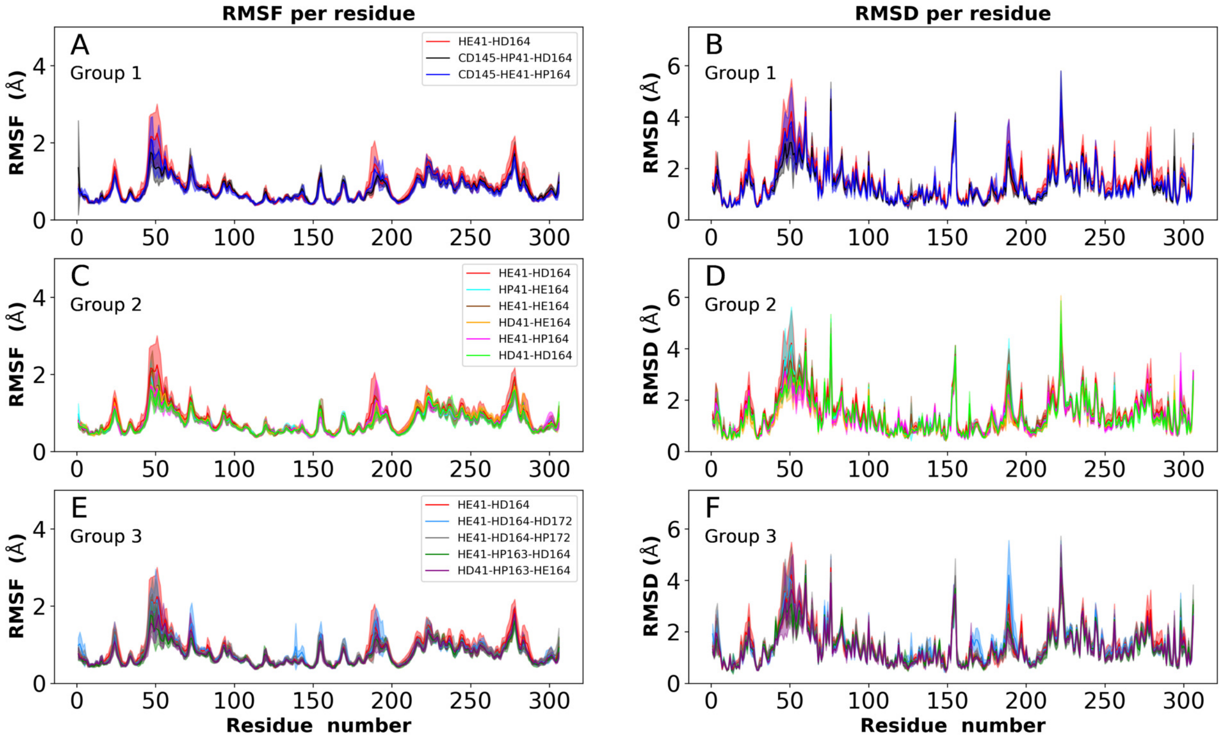


Figure S15: RMSF of $C_\alpha$ atoms (left) and RMSD of whole residues (right) for all simulations of the N3-bound structure.

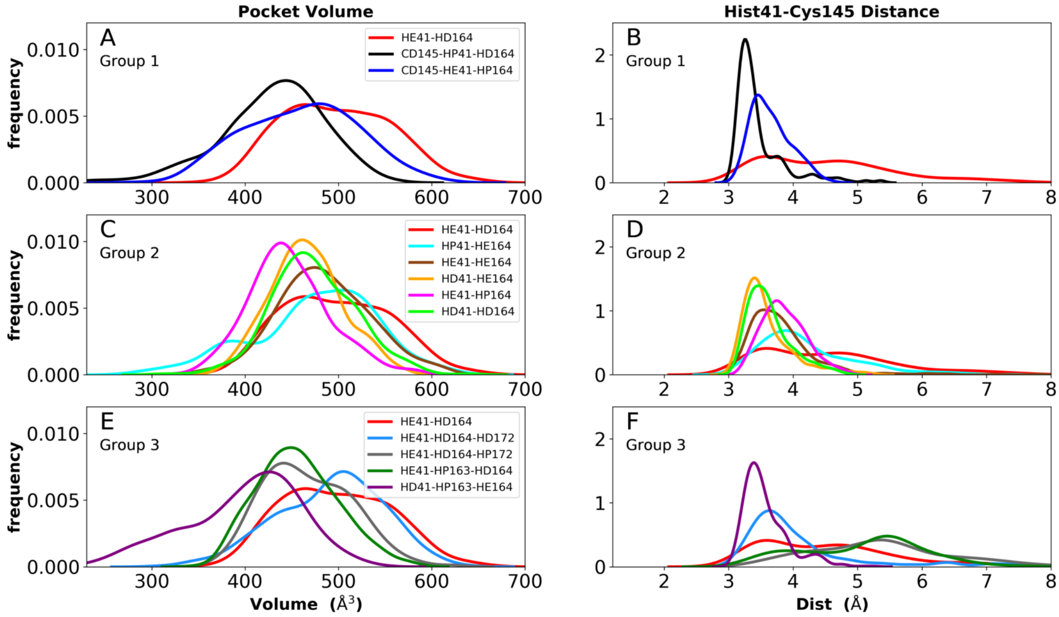


Figure S16: Distributions of pocket volume (left) and distance between NE and S in the catalytic residues His41 and Cys145 (right) for N3 bound structure.

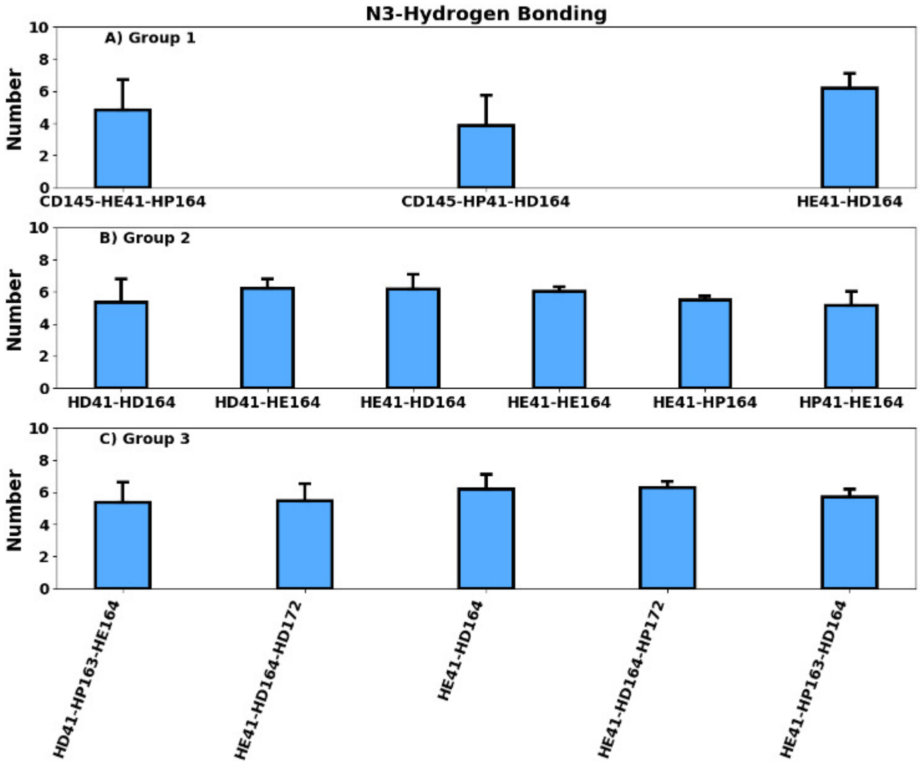


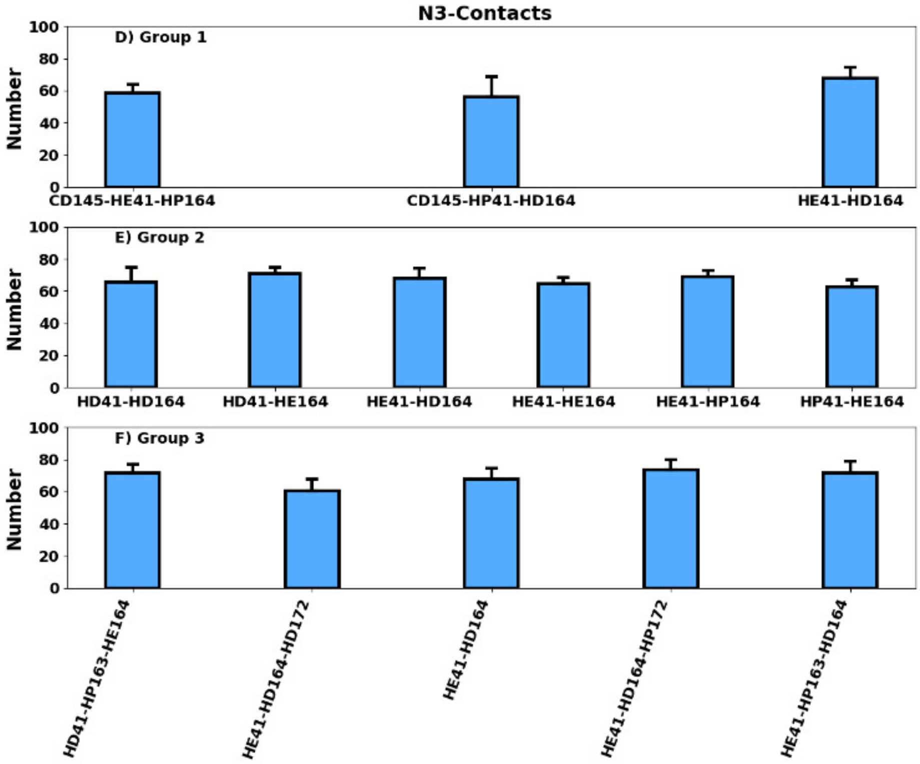


Figure S17: N3 hydrogen bonding and hydrophobic contacts. A-C) Total hydrogen bonds from the ligand to the protein and D-F) total hydrophobic contacts.

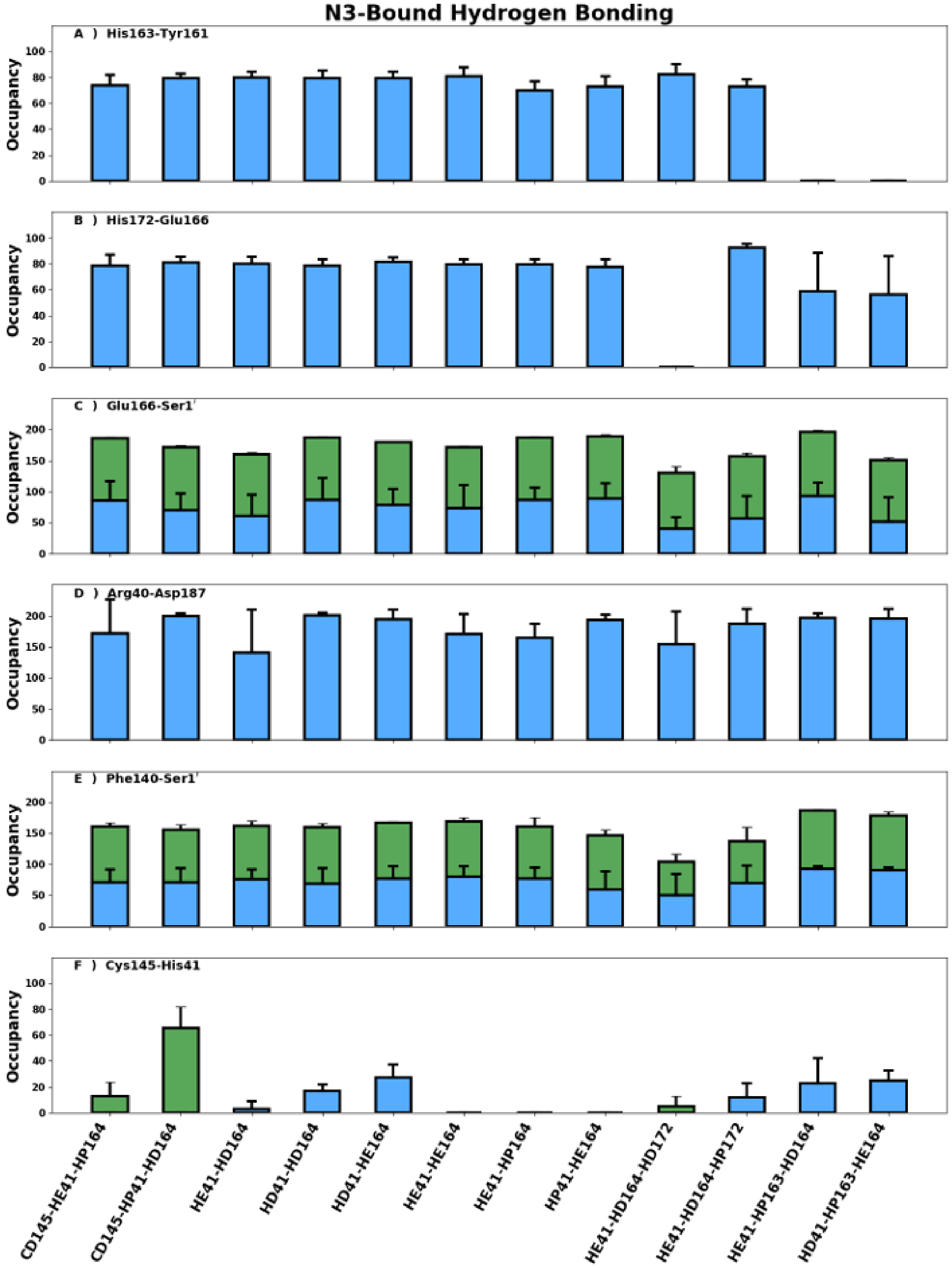


Figure S18: Hydrogen Bonding interactions in the N3-bound (PDB 7BQY) trajectories. A) His163-Tyr161, B) His172-Glu166, C) inter-monomer Glu166-Ser1′ with blue/green indicating the serine side chain/backbone, D) the Arg40-Asp187 salt bridge/charge reinforced hydrogen bond, E) inter-monomer interaction Phe140-Ser1′ with blue/green indicating Ser1′ acting as an acceptor/donor, and F) the catalytic dyad residues. In all figures, unless otherwise stated, His163 and His172 are HSE and standard deviations are indicated with bars. Also note, occupancies can be greater than 100% in cases where more than one hydrogen bond can be formed.

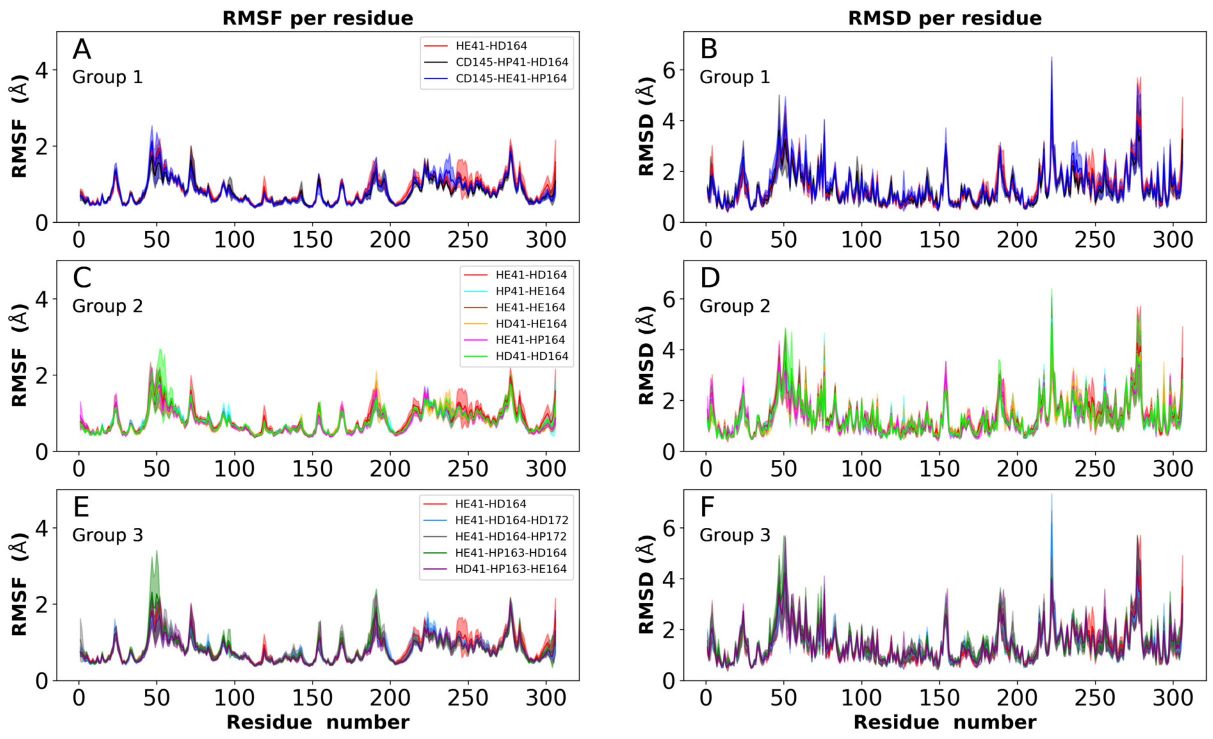


Figure S19: RMSF of $C_\alpha$ atoms (left) and RMSD of whole residues (right) for all simulations of ketoamide-bound structure 6Y2G.

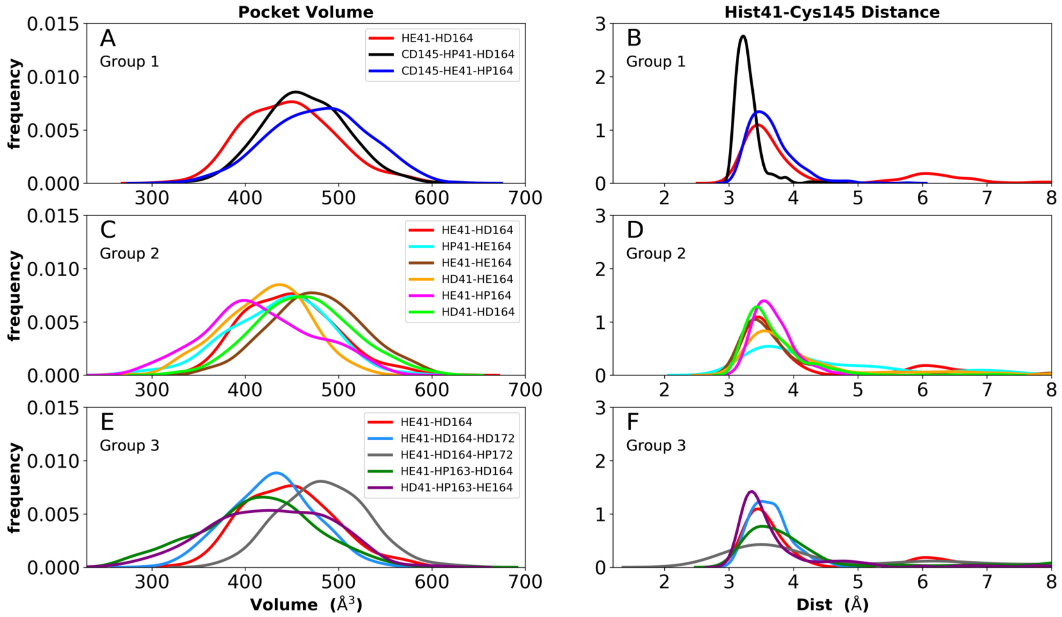


Figure S20: Distributions of pocket volume (left) and distance between NE and S in the catalytic residues His41 and Cys145 (right) for ketoamide-bound structure 6Y2G.

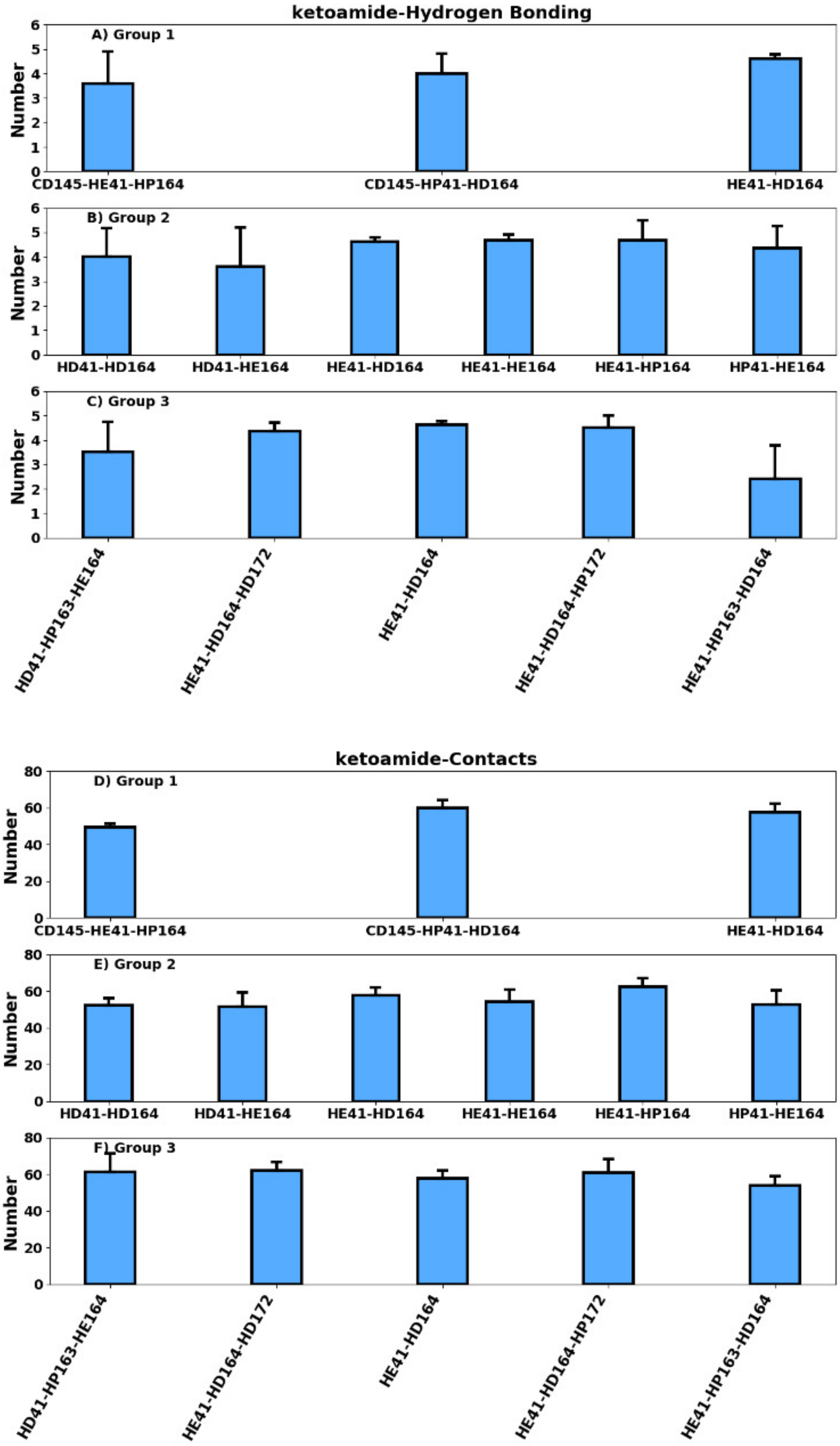


Figure S21: Ketoamide hydrogen bonding and hydrophobic contacts. A-C) Total hydrogen bonds from the ligand to the protein and D-F) total hydrophobic contacts.

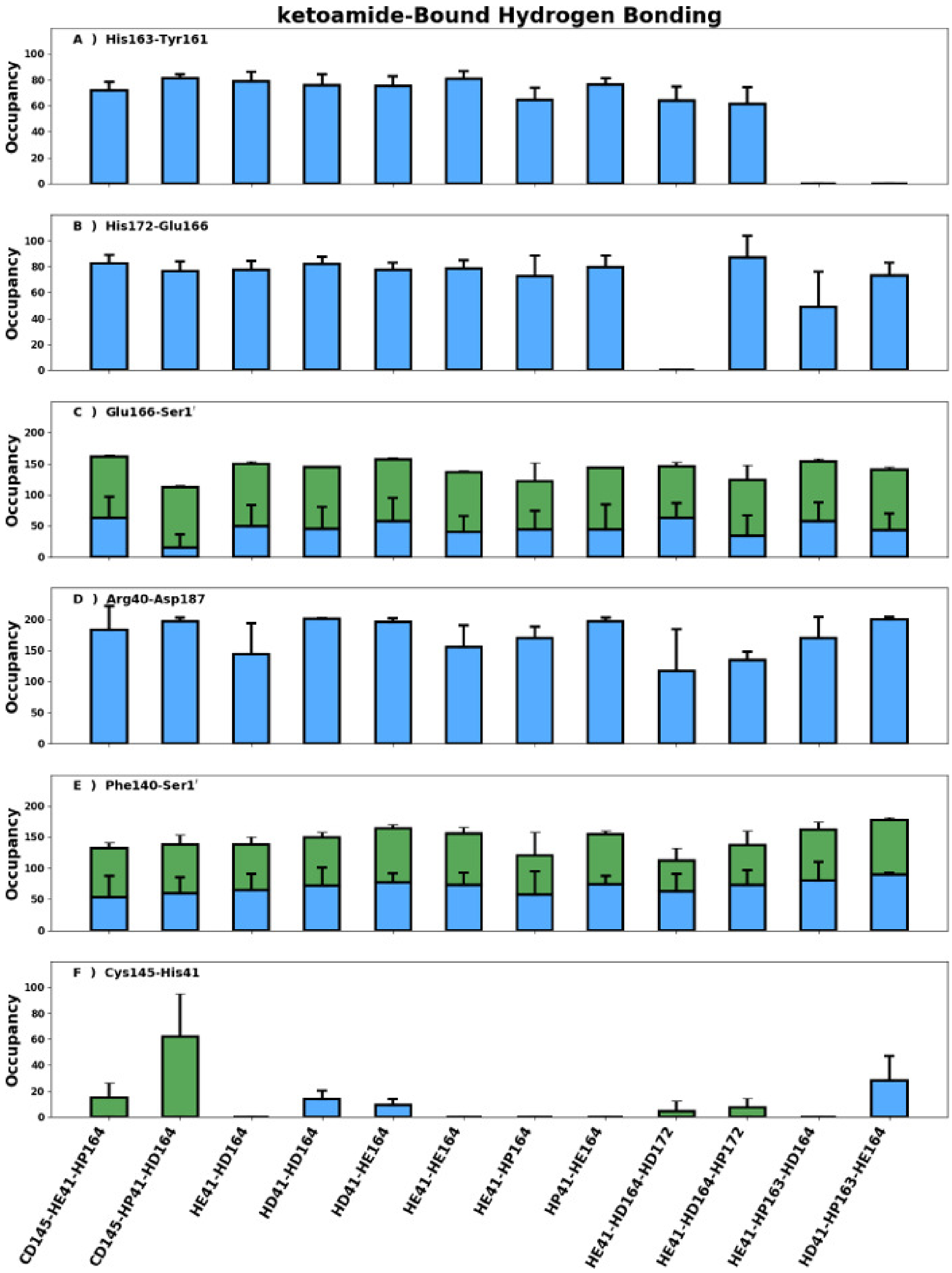


Figure S22: Hydrogen bonding interactions in the ketoamide-bound (PDB 6Y2G) trajectories. A) His163-Tyr161, B) His172-Glu166, C) inter-monomer Glu166-Ser1′ with blue/green indicating the serine side chain/backbone, D) the Arg40-Asp187 salt bridge/charge reinforced hydrogen bond, E) inter-monomer interaction Phe140-Ser1′ with blue/green indicating Ser1′ acting as an acceptor/donor, and F) the catalytic dyad residues. In all figures, unless otherwise stated, His163 and His172 are HSE and standard deviations are indicated with bars. Also note, occupancies can be greater than 100% in cases where more than one hydrogen bond can be formed.

Table S4: Individual free-energy perturbation simulations used to determine $\Delta\Delta G^\circ$ for the protonation state changes indicated in the N3-bound and ketoamide-bound states. Note that all numbers are for two concomitant transformations in the dimer. All energies are in kcal/mol.

| transformation | state | length | $\Delta G_{\text{forward}}$ | $\Delta G_{\text{backward}}$ | $\Delta G_{\text{BAR}}$ |
|---|---|---|---|---|---|
| HD41⟶HE41 | unbound | 160ns | -26.77 | +26.71 | -26.72±0.03 |
| HE163⟶HP163 | unbound | 160ns | +61.31 | -60.06 | +60.66±0.63 |
| HD41⟶HE41 | bound, ketoamide | 160ns | -29.07 | +29.74 | -29.40±0.34 |
| HE163⟶HP163 | bound, ketoamide | 160ns | +65.39 | -68.67 | +67.39±1.20 |
| HD41⟶HE41 | bound, N3 | 160ns | -25.32 | +25.09 | -25.19±0.12 |
| HE163⟶HP163 | bound, N3 | 160ns | +65.22 | -63.66 | +64.42±0.79 |
